# Mechanistic model of nutrient uptake explains dichotomy between marine oligotrophic and copiotrophic bacteria

**DOI:** 10.1101/2020.10.08.331785

**Authors:** Noele Norris, Naomi M. Levine, Vicente I. Fernandez, Roman Stocker

## Abstract

Marine heterotrophic bacteria use a spectrum of nutrient uptake strategies, from that of copiotrophs—which dominate in nutrient-rich environments—to that of oligotrophs—which dominate in nutrient-poor environments. While copiotrophs possess numerous phosphotransferase systems (PTS), oligotrophs lack PTS and rely on ATP-binding cassette (ABC) transporters, which use binding proteins. Here we present a molecular-level model that explains the dichotomy between oligotrophs and copiotrophs as the consequence of trade-offs between PTS and ABC transport. When we approximate ABC transport in Michaelis–Menten form, we find, contrary to the canonical formulation, that its half-saturation concentration *K*_M_ is not a constant but instead a function of binding protein abundance. Thus, oligotrophs can attain nanomolar *K*_M_ values using binding proteins with micromolar dissociation constants and while closely matching transport and metabolic capacities. However, this requires large periplasms and high abundances of binding proteins, whose slow diffusion limits uptake rate. We conclude that the use of binding proteins is critical for oligotrophic survival yet severely constrains maximal growth rates, thus fundamentally shaping the divergent evolution of oligotrophs and copiotrophs.

## Introduction

Approximately half of global carbon fixation occurs in the ocean (Falkowski *et al*, 1998). The fate of that carbon is governed by diverse species of heterotrophic bacteria (Giovannoni & Stingl, 2005; Azam & Malfatti, 2007; Burd *et al*, 2016) that differ in their carbon preferences and uptake rates (Poretsky *et al*, 2010; Gifford *et al*, 2013; Hellweger, 2018). Yet we lack a fundamental understanding of how and why species’ metabolic strategies differ, an understanding needed to predict how a changing climate will affect rates of carbon flux in the ocean (Widder *et al*, 2016).

An important driver of species’ differentiation is nutrient availability, leading to two divergent microbial lifestyles: copiotrophs dominate in nutrient-rich environments, whereas oligotrophs dominate in nutrient-poor environments (Koch, 2001; Giovannoni *et al*, 2014; Kirchman, 2016). A typical copiotroph exhibits a feast-and-famine lifestyle and swims to colonize sporadic, nutrient-rich patches and particles (Koch, 1971; Srinivasan & Kjelleberg, 1998). It reaches volumes greater than 1 μm^3^ and doubling times less than one hour (Lauro *et al*, 2009). Conversely, a typical oligotroph is nonmotile and free-living (Luo & Moran, 2015) and has volumes smaller than 0.1 μm^3^ and doubling times greater than 5 hours (Lauro *et al*, 2009). Although copiotrophs attain higher doubling rates and have larger per cell biomass, oligotrophs comprise the majority of marine bacterial biomass (Malmstrom *et al*, 2005; Yooseph *et al*, 2010). Despite this, most of our understanding of bacterial metabolism derives from research on copiotrophs, which are easier to culture (Gest, 2008).

Genomic analyses suggest that the divergent phenotypic traits of copiotrophs and oligotrophs are correlated with their suite of genes for nutrient transport (Lauro *et al*, 2009; Sowell *et al*, 2009; Cottrell & Kirchman, 2016; Noell & Giovannoni, 2019). Copiotrophs have many genes for phosphotransferase systems (PTS) used to uptake specific sugars (Kotrba *et al*, 2001). In contrast, oligotrophs lack PTS and instead rely heavily on ATP-binding cassette (ABC) transport systems, which are comprised of a transmembrane transport unit and a periplasmic substrate-binding protein. ABC transport systems have higher affinities than PTS (Neidhardt *et al*, 1990; Tam & Saier, 1993). Although it has long been held that the high affinity of ABC transport is a consequence of high-affinity binding proteins (Krupka, 1992; Bohl *et al*, 1995), it was recently suggested that ABC transport confers high affinity only when the abundance of binding proteins exceeds that of transport units (Bosdriesz *et al*, 2015). Thus, oligotrophs’ high abundances of binding proteins may explain their ability to grow in low nutrient conditions (Sowell *et al*, 2009; Giovannoni, 2017). However, it is not understood why oligotrophs cannot achieve higher grower rates in nutrient-rich conditions or why copiotrophs cannot achieve higher affinities in nutrient-limited conditions (Koch, 2001).

To understand the metabolic constraints governing oligotrophic versus copiotrophic lifestyles, we develop molecular-level transport and cellular proteome allocation models to compare the performance of ABC transport and PTS. We derive a Michaelis–Menten approximation of ABC transport kinetics, which shows, contrary to the classic paradigm, that the half-saturation concentration is not constant but varies with binding protein abundance. We thus find that ABC transport allows independent tuning of affinity and maximal uptake rate so that oligotrophs can achieve high affinities while closely matching transport and metabolic capacities. We predict that oligotrophs can attain a half-saturation concentration over a thousand-fold smaller than the binding protein’s dissociation constant. However, attaining this high affinity requires a great abundance of binding proteins, which, due to their large size, diffuse slowly and require large periplasms. Consequently, the reliance on binding proteins to achieve high affinity precludes high growth rates. Moreover, the ability of ABC transport to achieve high affinities while matching metabolic capacity makes metabolic imbalances unlikely and thus mechanisms for handling sudden nutrient up-shifts typically unnecessary, which may explain the toxicity of high-nutrient conditions to oligotrophs. Together, these findings provide the first mechanistic explanation for the divergence of copiotrophs and oligotrophs, as the consequence of trade-offs between PTS and ABC transport.

### Box 1

**Schematic of transport systems.**

For a nutrient to enter the cytoplasm, a transport unit bound to the inner membrane must expend energy to modify the substrate or translocate the substrate against a concentration gradient. For transport of a sugar by a phosphotransferase system (PTS), the sugar binds directly to the transport unit, and a cascade of specific proteins phosphorylate that particular sugar. For transport of a substrate by an ATP-binding cassette (ABC) transport system, binding proteins in the periplasm first scavenge for and store the substrate in the periplasm. When bound to substrate, a binding protein can then bind to a membrane-bound transport unit, which uses ATP to translocate the substrate. While a single type of binding protein may be able to bind to different substrates, it can bind to only a single, corresponding type of transport unit. To limit the number of free parameters when modeling these two transport systems, we use a simple model of PTS that assumes that binding of the substrate to the transport unit is irreversible. We extend the model for ABC transport to account for the reversible binding of the substrate to the binding protein.

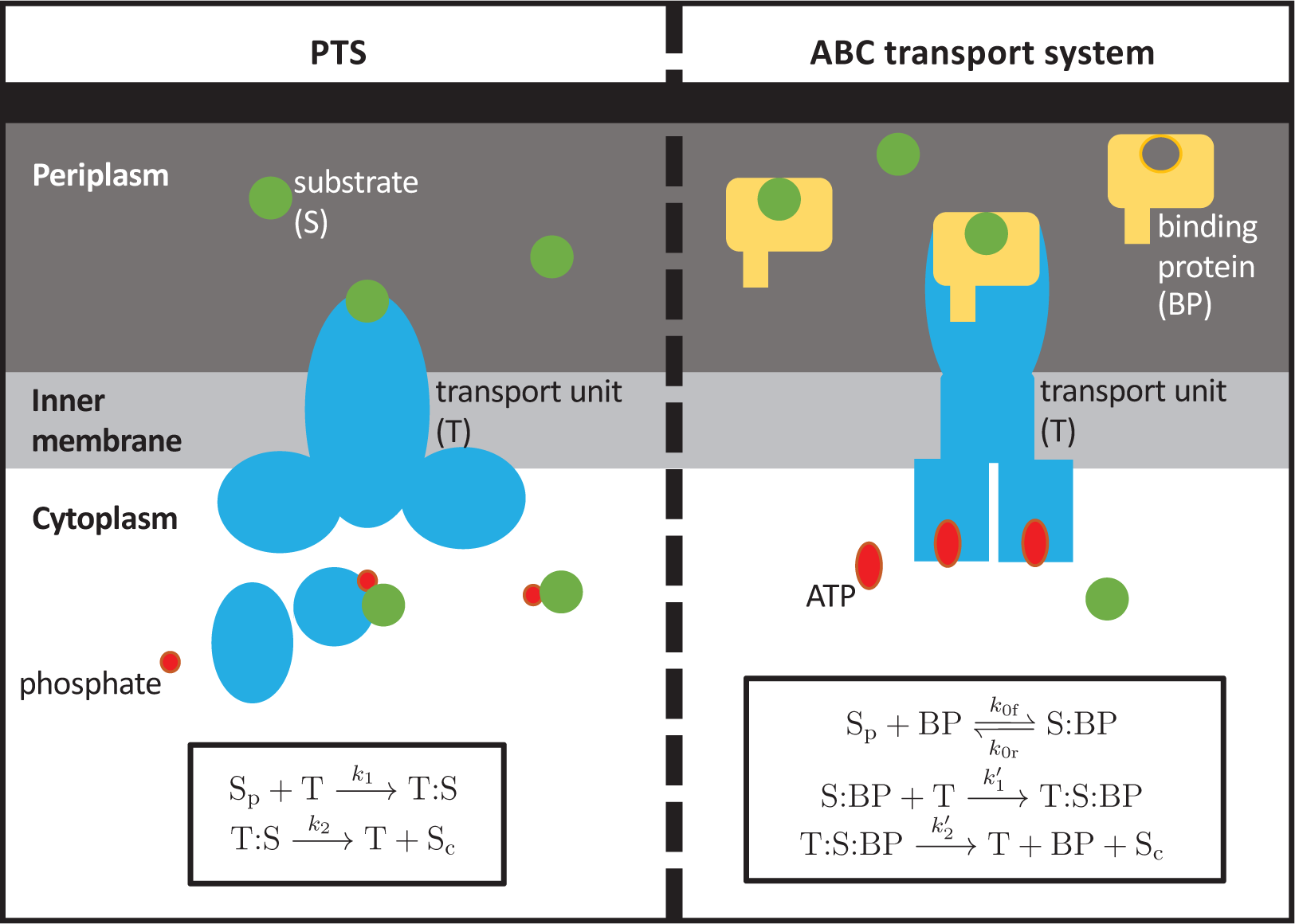

## Results

### The half-saturation concentration of ABC Michaelis–Menten kinetics is not constant

To contrast the nutrient acquisition strategies of PTS and ABC transport systems, we derive models of both, which show that, whereas the half-saturation concentration of PTS is an intrinsic property of the transporter, the half-saturation concentration of ABC transport is a function of binding protein abundance. A PTS is used for the cytoplasmic uptake of a specific sugar and modifies the sugar once it enters the cytoplasm by binding the sugar to a phosphate group (Romano & Saler, 1992; Saler & Reizer, 1994; Siebold *et al*, 2001; Kotrba *et al*, 2001). PTS uptake kinetics can be described by the canonical model for transport (Lehninger *et al*, 2005). It describes transport as a two-step reaction, in which (*i*) the periplasmic substrate (S_p_) binds to the membrane-bound transport unit (T) with rate constant *k*_1_ to form a bound complex (T: S), and (*ii*) the substrate is translocated irreversibly into the cytoplasm with rate *k*_2_ (S_c_) (Box 1, Supplemental Section 1). Using mass-action kinetics, we find that the cytoplasmic uptake rate (the rate at which S_p_ is converted to S_c_) for PTS at steady-state is

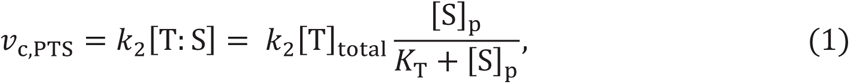

where *K*_T_ = *k*_2_/*k*_1_ is the transport unit dissociation constant and [T]_total_ is the concentration of membrane-bound transport units in the periplasm. (Note that we express all transport rates in terms of change in periplasmic concentration.) The solution in Equation 1 has the classic Michaelis–Menten form of nutrient transport (Michaelis *et al*, 2011), with maximal uptake rate *V*_max_ proportional to [T]_total_ and half-saturation constant *K*_M_ equal to *K*_T_ (Box 2).

In contrast to PTS, the kinetics of ABC transport does not follow the classic Michaelis–Menten form (Bosdriesz *et al*, 2015). ABC transport uses binding proteins (BP) in the periplasm that scavenge for the incoming nutrients. These binding proteins, when in complex with the substrate, bind to membrane-bound transport units that require ATP to translocate the substrate from the periplasm into the cytoplasm (Hengge & Boos, 1983; Hosie & Poole, 2001; Davidson *et al*, 2008). We thus describe ABC uptake by extending the PTS model to account for a three-step reaction (Bohl *et al*, 1995; Bosdriesz *et al*, 2015): (*i*) the substrate–binding protein complex (S: BP) is formed by a reversible reaction with association rate *k*_0f_ and dissociation rate *k*_0r_,(*ii*) the bound complex of substrate and binding protein (S: BP) binds with rate constant *k*^′^_1_ to the membrane-bound transport unit (T) to form a bound complex (T: S: BP), and (*iii*) the substrate is translocated irreversibly into the cytoplasm (S_c_) with rate *k*^′^_2_ (Box 1). At steady state, we obtain a system of four equations that can be solved exactly for the cytoplasmic uptake rate for ABC transport, *v*_c,ABC_, as a function of the concentration of free substrate in the periplasm, [S]_p_ (Supplemental Sections 1.2 and 4.3.2):

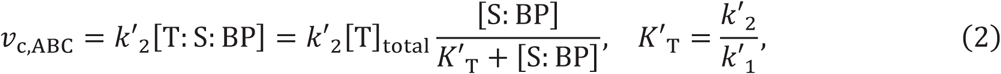

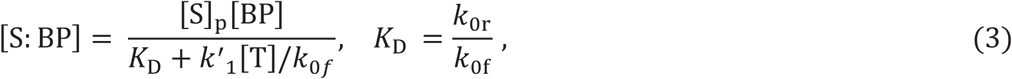

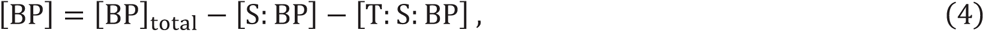

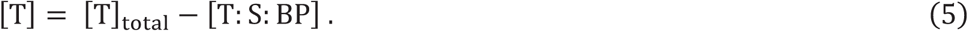

We used previous experimental observations of the well-characterized maltose ABC transport system in *Escherichia coli* to substantiate this model (Supplemental Section 3). The model accurately predicts the observed *K*_M_ as well as the shape of the uptake rate curves as functions of both extracellular maltose concentration and binding protein abundance (Supplementary Figures S1-S4).

#### Box 2

**Comparison of maximal uptake rates, half-saturation concentrations, and specific affinities for PTS and ABC transport systems.**

We can approximate cytoplasmic uptake rates using the Michaelis–Menten equation: *v*_c_ = *V*_max_[S]_p_/(*K*_M_ + [S]_p_), where *V*_max_ is the maximal uptake rate and *K*_M_ the half-saturation concentration. While the exact solution of the cytoplasmic uptake rate for our model of PTS transport is in the form of a Michaelis–Menten equation, the exact solution of the uptake rate for ABC transport is not. Because our simulations suggest that the abundance of binding proteins must exceed the abundance of transport units, we make the approximations that (*i*) [T: S: BP] ≪ [BP]_total_ and (*ii*) *k*^′^_1_[T] ≪ *k*_0r_ (Supplemental Section 2) to obtain the above estimates for the effective maximal rate and half-saturation concentration. For PTS, the half-saturation concentration is a constant equal to the dissociation constant *K*_T_ = *k*_2_/*k*_1_. For ABC transport, the half-saturation concentration depends on both the transport dissociation constant *K*^′^_T_ = *k*^′^_2_/*k*^′^_1_ and the binding protein dissociation constant *K*_D_ = *k*_0r_/*k*_0f_ and is additionally a function of the abundance of binding proteins. The specific affinity *a* = *V*_max_/*K*_M_ of ABC transport is thus proportional to the product of the abundances of transport units and of binding proteins.

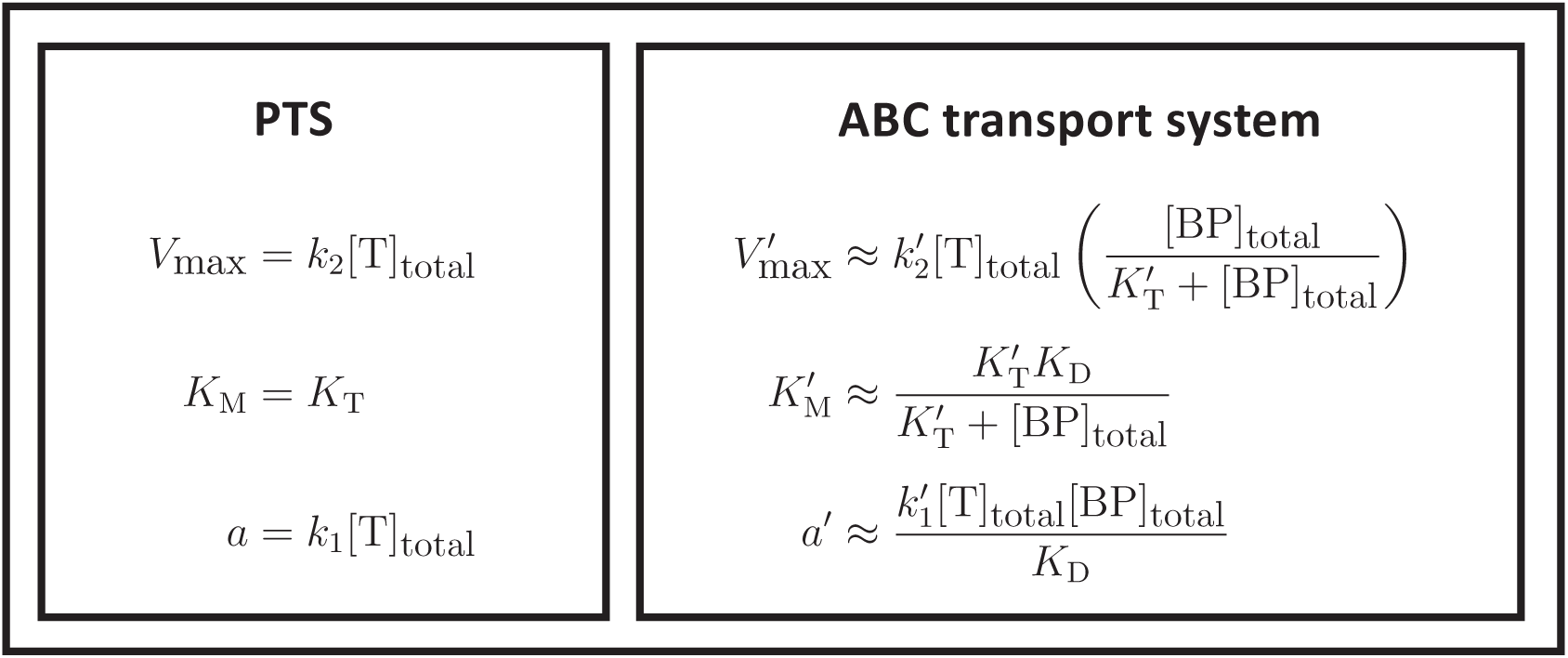

To obtain a compact analytical expression describing how transport protein abundance affects uptake rate, we derived an approximation of ABC transport kinetics in Michaelis–Menten form. By assuming that binding proteins are much more abundant than active transport units (Bosdriesz *et al*, 2015; Peebo *et al*, 2015) ([BP]_total_ ≫ [T: S: BP]) and that the abundance of inactive transport units is low (so that [T] ≪ *k*_0r_/*k*^′^_1_) (Box 1, Supplemental Section 2), we obtain from Equations 2–5 the following approximation for the cytoplasmic uptake rate:

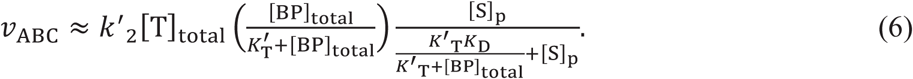

This Michaelis–Menten approximation captures the dynamics of the full ABC transport model (Equations 2–5) over a wide range of parameter values (Supplementary Figure S6).

This formulation clearly shows that the half-saturation “constant” *K*_M_ is, in fact, not a constant for ABC transport. This finding is in direct contrast to the classic and ubiquitously used Michaelis–Menten formulation of transport and to the traditional assumption that the value of *K*_M_ for ABC transport is a constant that is intrinsic to the binding proteins (Button, 1998; Noell & Giovannoni, 2019). Instead, we find that *K*_M_ depends on both the dissociation constant of the substrate and binding protein (*K*_D_) and the dissociation constant of the resulting complex and the transport unit (*K*^′^_T_). Importantly, we find that *K*_M_ is also a function of the concentration of binding proteins in the periplasm ([BP]_total_) (Box 2). Therefore, expressing high abundances of binding proteins enables oligotrophs to attain small *K*_M_ values and thus high affinities. At low nutrient concentrations, the classic Michaelis–Menten uptake rate is proportional to the specific affinity (Button *et al*, 2004), *a* = *V*_max_/*K*_M_. Whereas a bacterium using PTS has constant *K*_M_ and thus can increase its specific affinity in oligotrophic conditions only by tuning *V*_max_ (via expression of the transport unit; Equation 1), a bacterium using ABC transport can increase its specific affinity by tuning either *V*_max_ or *K*_M_ by tuning the expression levels of the transport units and binding proteins, respectively (Equation 6).

### A rate–affinity trade-off drives the differentiation of oligotrophs and copiotrophs

The derived Michaelis–Menten kinetics (Equations 1 and 6) suggest that ABC transport systems allow bacteria to achieve higher substrate affinities than PTS by expressing high abundances of binding proteins. To understand the costs associated with achieving these high affinities and thus to determine how the optimal expression levels of transport units and binding proteins differ in low-nutrient and high-nutrient environments, we integrate our solutions for the cytoplasmic uptake rates of PTS (Equation 1) and ABC transport (Equations 2–5) into a mechanistic, single-cell metabolic model (Figure 1, Supplementary Section 4). Similar to the self-replicator model (Molenaar *et al*, 2009), our metabolic model accounts for four protein groups – transport proteins, metabolic proteins, ribosomes, and membrane biosynthesis proteins – and is used to solve a proteome allocation problem that determines the optimal amount of each protein group that the cell should express in order to maximize its growth rate for a given extracellular nutrient concentration.

**Figure 1:**
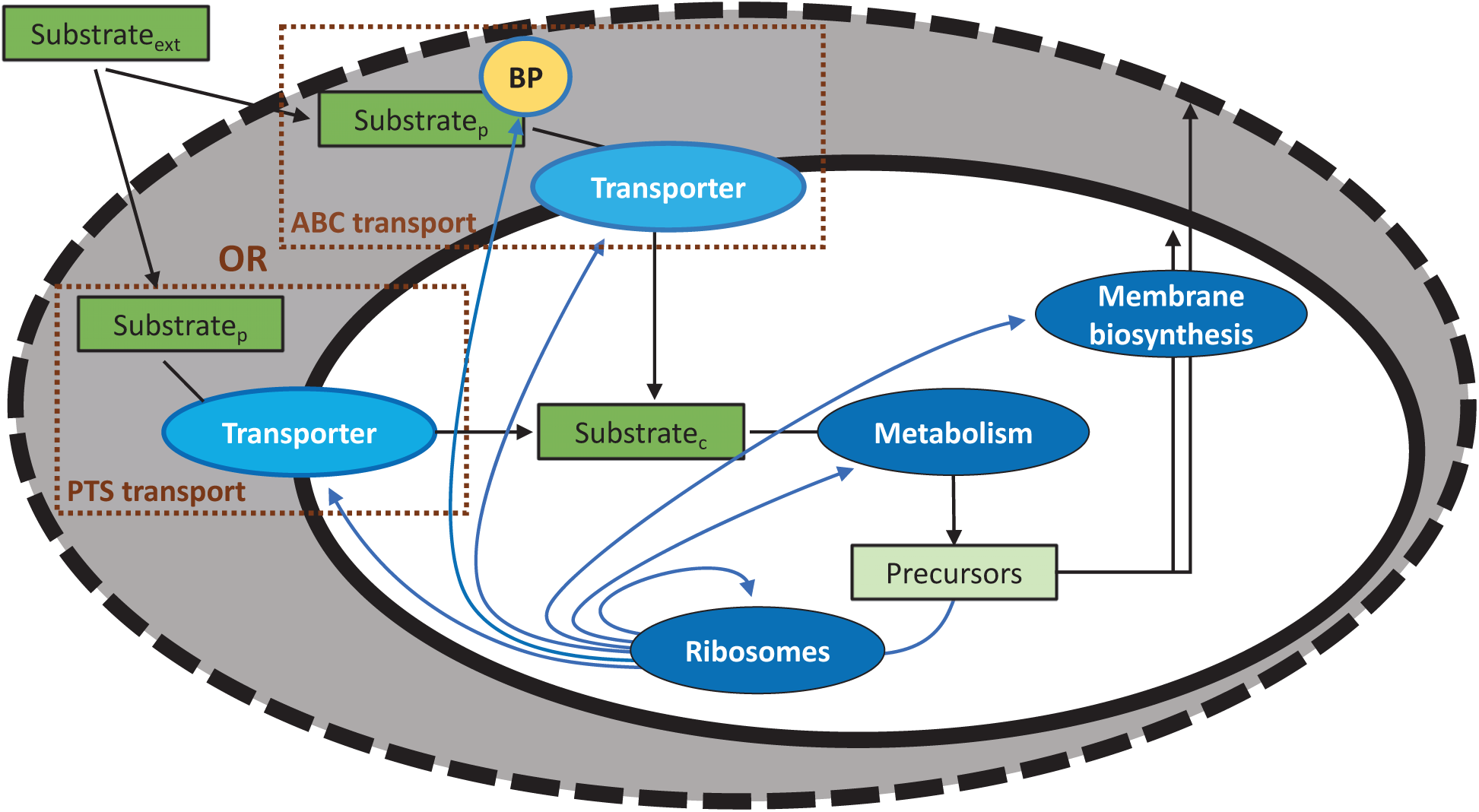
A simple metabolic model tracks the utilization of a generic nutrient by the cell. The nutrient diffuses into the periplasm via a porous outer membrane and is then transported into the cytoplasm by membrane-bound transport units. The cell uses either transport by PTS, in which the substrate directly binds to the transport unit, or ABC transport, in which the substrate must first bind to a binding protein and then this complex binds to the transport unit. The intracellular substrate is next metabolized by a protein group that transforms the substrate into a precursor (a generic amino acid) that is needed to build the cell. The precursors are used (*i*) by a membrane biosynthesis protein group to build both the outer and inner membranes and (*ii*) by ribosomes to make proteins comprising the six protein groups. This model is subject to a number of constraints to determine the proteome allocation that maximizes the steady-state exponential growth rate. While this model does not consider the utilization of carbon for energy, we expanded the model to consider energy to show that differences in the energetic requirements of PTS and ABC transport do not change our results (Supplemental Section 5).

Our metabolic model tracks the transport of a nutrient into the cytoplasm and the subsequent transformation of that nutrient into the proteins and metabolites required for replication. The abundances of proteins and metabolites are constrained by the cell’s surface-area-to-volume ratio. Because the cellular components occupy volume, they are limited by maximum cytoplasmic and periplasmic densities (Molenaar *et al*, 2009), which favors smaller surface-area-to-volume ratios. On the other hand, the surface of the inner membrane must be sufficiently large because the membrane-bound transport units carry “real estate costs” (Molenaar *et al*, 2009; Szenk *et al*, 2017). Larger surface-area-to-volume ratios also support higher specific uptake rates by diffusion at low-nutrient conditions (Koch, 1996; Young, 2006; Armstrong, 2008). Thus, taken together, the surface-area-to-volume ratio creates a trade-off between the cell’s capacity for uptake and its capacity for synthesis. Therefore, in addition to determining the optimal proteome allocations, our model also determines the optimal surface-area-to-volume ratio, the protein and metabolite concentrations that are constrained by this ratio, and the fraction of the volume devoted to the periplasm (Supplemental Section 4).

Central to this optimization problem are the costs and benefits of expressing more of a particular protein group. While expressing more transport units or binding proteins increases the uptake rate, it incurs a proteomic cost (Kaleta *et al*, 2013; Peebo *et al*, 2015; Noor *et al*, 2016; Basan, 2018). This cost is an opportunity cost. For example, because growth rate depends on the proteome fraction allocated to ribosomes (Scott *et al*, 2014), expressing greater abundances of transport proteins may limit growth, as it limits the proportion of the proteome devoted to ribosomes. We assume that the transport units of PTS and ABC transport systems have the same proteomic cost and that the proteomic cost of an ABC binding protein is four times less than the cost of a transport unit (see Supplemental Section 4.3 for justifications).

The effect that these transport proteomic costs have on the optimal proteome allocation strongly depends on the uptake rate per transport unit. This uptake rate is often limited by rates of diffusion within the periplasm (Brass *et al*, 1986). Hence, we argue that differences in substrate diffusion across the periplasm drive a trade-off between PTS and ABC transport. Because ABC binding proteins are much larger than the substrates they bind, the diffusivity of the binding proteins is lower than the diffusivity of the substrate, limiting the achievable rates of ABC transport relative to PTS (Bosdriesz *et al*, 2015). For example, a typical binding protein (MalE) has a molecular weight of approximately 40 kDa and thus an estimated cytoplasmic diffusivity of 2 μm^2^/s, whereas glucose has a molecular weight of 0.18 kDa and thus an estimated cytoplasmic diffusivity of 200 μm^2^/s (Trovato & Tozzini, 2014). Because the periplasmic diffusivities of the binding proteins and the kinetic rates of ABC transport systems are unknown (Rees *et al*, 2009), our model therefore assumes that the association rate *k*^′^_2_ and the translocation rate *k*^′^_1_ (Box 1) are one-hundred times smaller than the equivalent rates for PTS (i.e., *k*^′^_1_ = 0.01*k*_1_, *k*^′^_2_ = 0.01*k*_2_) because both rates depend on diffusion. Sensitivity analyses of all transport kinetics parameters confirmed that the fundamental trends emerging from our results do not depend on the precise values chosen (Supplementary Figures S9, S10).

Because the diffusive rates of binding proteins limit ABC uptake rates, our model shows that PTS can achieve higher maximal uptake rates *V*_max_ per proteomic cost than ABC transport (Box 2, Supplementary Figure S7*C*). Specifically, at saturating extracellular nutrient concentrations, the optimal cell using PTS devotes 80 times less proteome to transport than the optimal cell using ABC transport (Figure 3*A*) yet achieves a slightly (3%) higher *V*_max._ (Supplementary Figure S8*A*). Therefore, cells using PTS achieve higher growth rates than cells using ABC transport when nutrient concentrations are high (Figure 2*A*).

**Figure 2:**
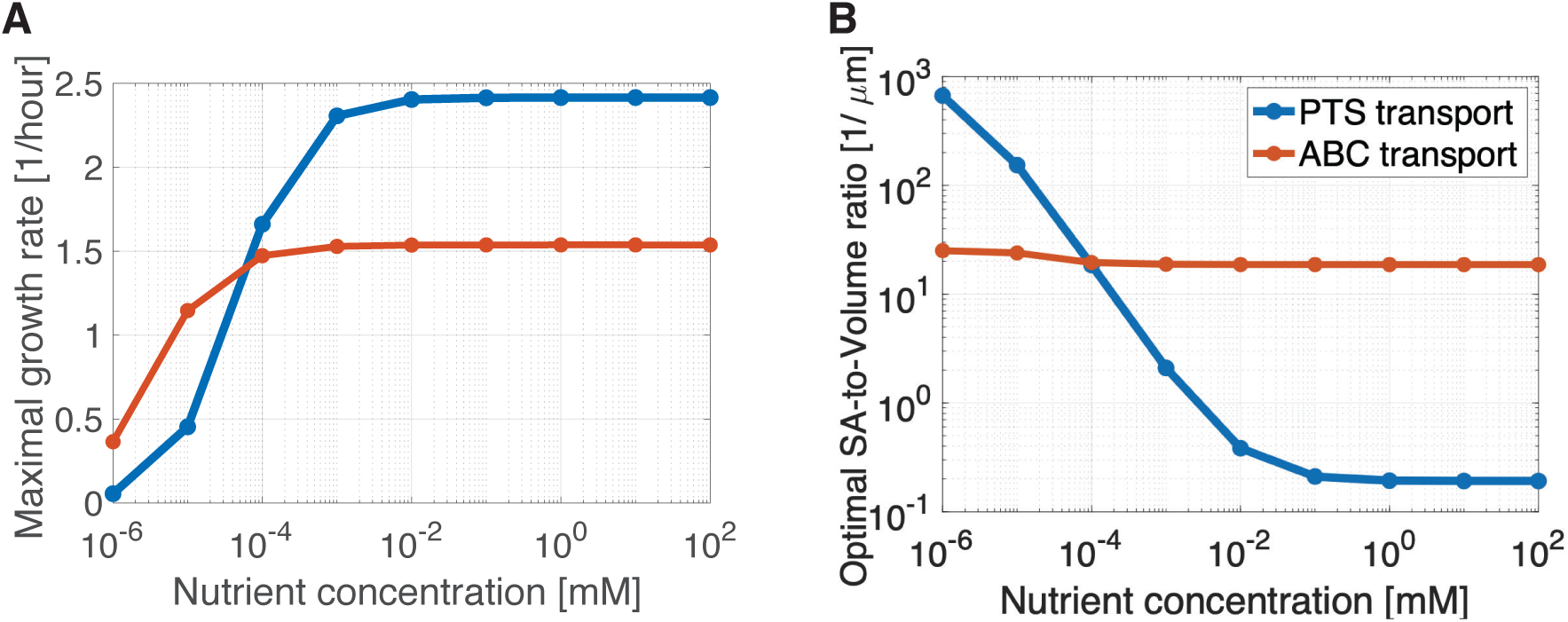
A rate–affinity trade-off. Plots show the results of proteome allocation problems using either PTS or ABC transport and solved for different extracellular nutrient concentrations (x-axis). We assume that the association and translocation rates are 100 times lower for ABC transport than for PTS (*k*^′^_1_ = 0.01*k*_1_, *k*^′^_2_ = 0.01*k*.). (A) shows the maximal growth rates achieved using the optimal proteome allocation, and (B) shows the optimal surface-area-to-volume ratio used to achieve those maximal growth rates. As can also be seen in Figures S7&S8, ABC transport achieves higher growth rates at low nutrient concentrations because they support higher substrate affinities per transport proteomic cost, whereas PTS achieves higher growth rates at high nutrient concentrations because they support higher maximal uptake rates per transport proteomic cost.

**Figure 3:**
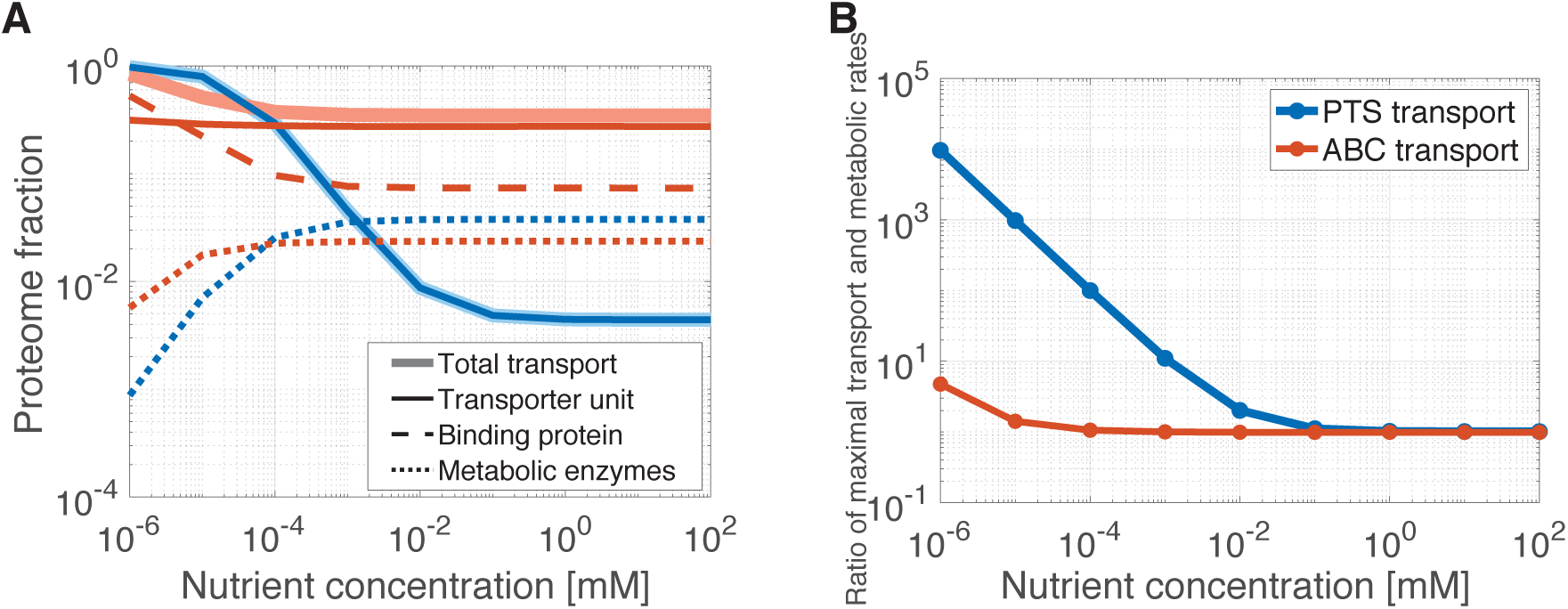
Optimal proteome allocation for PTS and ABC transport systems. (A) While it is optimal for cells relying on either PTS or ABC transport systems to devote nearly all of their proteome to transport at low nutrient concentrations, for ABC transport systems, it is the proteome fraction of the binding proteins that increases as nutrient concentration decreases and not the fraction allocated to the membrane-bound transport units. (B) As the nutrient concentration decreases, the optimal maximal uptake of transport increases for PTS but decreases for ABC transport systems. This results in an increasing ratio of optimal maximal uptake and maximal metabolic rates for transport by PTS as nutrient concentration decreases, while it is optimal for ABC transport systems to maintain this ratio closer to one.

Conversely, our model shows that ABC transport systems have higher specific affinities (*a*) per proteomic cost than PTS (Box 2, Supplementary Figure S7*C*). At 1 nM extracellular nutrient concentration, the optimal ABC cell has a binding protein to transport unit ratio ([BP]/[T]_total_) of seven (Figure 4). In this way, using a binding protein with *K*_D_ = 1 *μ*M and a transport unit with dissociation constant *K*^′^_T_= 10 *μ*M, the cell achieves an effective half-saturation concentration of *K*^′^_M_ = 3 nM (Figure 4). Thus, although the optimal ABC cell devotes 16% less of its proteome to transport than the optimal PTS cell, it achieves a half-saturation concentration that is over three thousand times lower than the half-saturation concentration of the optimal PTS cell, *K*_M_ = *K*_T_ = 10 *μ*M. Therefore, cells using ABC transport achieve higher growth rates than cells using PTS when nutrient concentrations are low (Figure 2*A*).

**Figure 4:**
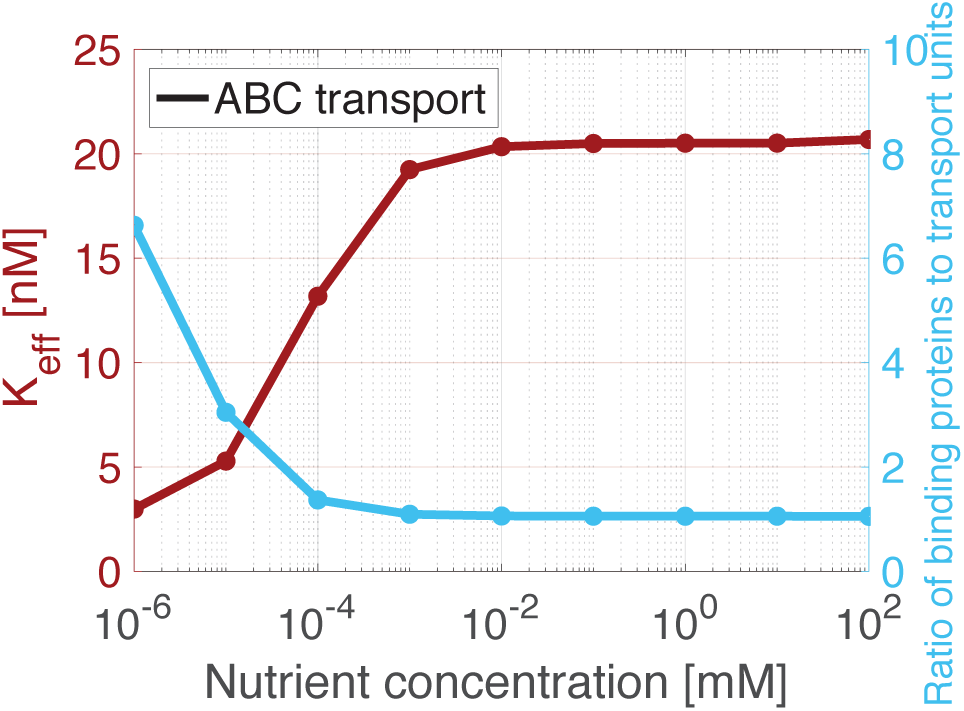
The effective half-saturation concentration of ABC transport. ABC transport systems achieve low optimal half-saturation concentrations (*K*_eff_, magenta curve and axis)—and thus high specific affinities—as nutrient concentrations decrease by maintaining a high surface-area-to-volume ratio and increasing the ratio of the abundance of binding proteins to the abundance of membrane-bound transport units (turquoise curve and axis). (For a plot showing how we calculate the effective half-saturation concentration, see Figure S11.)

Many bacterial species have both PTS and ABC transport systems for the same nutrient, using PTS when the nutrient is plentiful and ABC transport when the nutrient is scarce (Neidhardt *et al*, 1990; Schleif, 2010). Because of this redundancy, it has long been hypothesized that there exists a rate-affinity trade-off between PTS and ABC transport (Neidhardt *et al*, 1990; Bosdriesz *et al*, 2015). Our results provide a mechanistic explanation for this trade-off and furthermore demonstrate that this trade-off, in particular, drives the differences in performance between the optimal ABC and PTS cell. Alternative hypotheses on the mechanisms creating a trade-off between the two transport mechanisms are not supported by our metabolic model. We find that the advantage of PTS in high-nutrient conditions does not stem from either lower energetic or lower proteomic costs because these costs are minimal in high-nutrient conditions. Specifically, when we expanded our model to include the energetic costs of transport and furthermore incorrectly assumed that the association rate, *k*_1_, and the translocation rate, *k*_2_, were the same for both PTS and ABC transport systems, we observed no trade-off: despite the higher proteomic and energetic costs of ABC transport, the maximal growth rate achieved by the optimal ABC cell was always greater than or equal to the maximal growth rate achieved by the optimal PTS cell (Supplementary Section 5; Supplementary Figure S12*A*). We therefore conclude that the only trade-off that can explain the redundancy of species that utilize both PTS and ABC transport systems for the same nutrient is a rate–affinity trade-off that is a consequence of the high affinity achieved by using binding proteins and the binding proteins’ limiting rates of diffusion. Oligotrophs’ lack of PTS and heavy reliance on ABC transport suggest that this trade-off is central to the dichotomy between copiotrophs and oligotrophs and that it may explain the fundamental difference in their achievable growth rates.

Our conclusion that a rate–affinity trade-off between PTS and ABC transport underpins the differentiation of oligotrophs and copiotrophs is further supported by our model’s predictions on the optimal surface-area-to-volume ratio. We find that the optimal surface-area-to-volume ratio (which is inversely proportional to the cell radius) is smaller for PTS cells in nutrient-rich conditions than it is for PTS cells in nutrient-poor conditions or for ABC cells in all nutrient conditions (Figure 2*B*). This result is in accordance with observations that copiotrophic marine bacteria are typically over ten times larger than oligotrophic ones (Young, 2006; Lauro *et al*, 2009). The model further reveals that increasing transport translocation rates decreases the optimal surface-area-to-volume ratio (Supplementary Figures S13, S14). This indicates that, whereas larger surface-area-to-volume ratios allow the cell to achieve higher cytoplasmic concentrations of metabolites and proteins—and hence higher processing rates—in low-nutrient conditions, smaller surface-area-to-volume ratios are optimal at high nutrient uptake rates because they provide the cell with more space in which to process the substrate and transform it into biomass. Thus, the higher achievable uptake rates of PTS support larger optimal cell volumes.

### Cells relying on ABC transport can achieve high affinities while closely matching their metabolic and transport capacities

Unlike a cell using PTS, a cell using ABC transport can increase its specific affinity without increasing its maximal uptake rate (Equation 6). Our model predicts that, for both the optimal PTS cell and the optimal ABC cell, as the extracellular nutrient concentration decreases, the fraction of the proteome devoted to transport increases, while the fraction of the proteome devoted to metabolic enzymes decreases (Figure 3*A*). For PTS, increasing the transport proteome fraction increases the maximal uptake rate *V*_max_ (Box 2). As a result, our model demonstrates a mismatch between metabolic and transport capacities for a PTS cell optimized for growth in low-nutrient conditions: for the optimal PTS cell, the ratio of the transport capacity *V*_max_ and the metabolic capacity—which is proportional to the abundance of metabolic enzymes—exceeds one for all nutrient concentrations below 1 mM. Indeed, at 1 nM, this ratio approaches 10,000 (Figure 3*B*). Therefore, a PTS cell optimized for growth in nutrient-poor conditions that suddenly encounters a higher nutrient concentration would uptake more nutrient than it can process and could quickly accumulate toxic levels of metabolites if it cannot excrete them.

In contrast, as the extracellular nutrient concentration decreases, the optimal ABC cell does not allocate any additional proteome to membrane-bound transport units but only to binding proteins to increase its affinity (Figure 3*A*). As a result, our model shows that the optimal ABC cell maintains a transport to metabolic capacity ratio of one for all nutrient concentrations above 0.1 μM. At 1 nM, the maximum ratio is below ten (Figure 3*B*). Therefore, our model suggests that a cell using ABC transport is much less prone to mismatches between its proteome and the environment that may cause toxic accumulations of metabolites within the cell. Hence, oligotrophs may never need to excrete metabolites.

## Discussion

We used a metabolic model to quantify the costs and benefits of using PTS versus ABC transport systems and thus understand the divergence of the copiotrophic and oligotrophic lifestyles of marine bacteria. By deriving an approximation of ABC transport in Michaelis–Menten form, we found that we must reconsider the canonical representation of transport, which assumes a constant half-saturation concentration (Lehninger *et al*, 2005; Michaelis *et al*, 2011). Our approximation shows how the half-saturation concentration *K*_M_ varies with binding protein abundance, corroborating previous theoretical work that found that attaining low *K*_M_ values requires binding protein abundances in excess of transport unit abundances (Bosdriesz *et al*, 2015). Furthermore, we predict that, for oligotrophic bacteria, the *K*_M_ value may be over a thousand-fold smaller than the dissociation constant of the binding protein *K*_D_. Although we are aware of only two experimental studies that considered the effects of varying binding protein abundance on uptake, both provide support to our model. We used one of the experimental studies—on *E. coli*’s ABC maltose transport system—to directly verify our model’s predictions on the dependence of uptake on binding protein abundance (Supplementary Section 3). The second study found that a *Salmonella typhimurium* mutant that expresses fivefold higher levels of binding protein for histidine uptake has a fourfold lower *K*_M_—thirtyfold lower than the estimated *K*_D_ of the binding protein for histidine (Ames & Lever, 1970).

As ABC transport systems are ubiquitous in gram-negative bacteria, the fact that *K*_m_ may be drastically different from *K*_D_ has important implications for our ability to predict microbial dynamics. Because of the difficulty of measuring the value of *K*_M_ for uptake directly, much previous work has estimated the performance of ABC transport systems using binding assays that measure *K*_D_ instead (Tam & Saier, 1993; Davidson *et al*, 2008). Our work suggests that this estimate could differ from *K*_M_ by orders of magnitude for oligotrophs that use high abundances of binding proteins, thus potentially leading to substantial underestimates of oligotrophs’ nutrient uptake rates. In addition, a variety of microbial ecosystem models assume a constant value of *K*_M_ for uptake (Tilman, 1977; Zakem & Levine, 2019), but this assumption may be flawed because bacteria may vary their binding protein abundance and thus their *K*_M_ value as a function of environmental conditions. Alternatively, it is also possible that cells have evolved to express a constant binding protein abundance to maintain a constant, ecologically relevant *K*_M_ value. Experiments are needed to determine the extent to which binding protein abundance and the value of *K*_M_ vary within a species, as well as the impacts of the variability in *K*_M_ on ecosystem dynamics.

Our model provides a mechanistic explanation for a seemingly puzzling discrepancy between the glycine betaine transport systems of an oligotrophic marine bacterium of the SAR11 clade and *E. coli*. The SAR11 strain *Pelagibacter ubique* can achieve nanomolar values for the half-saturation concentration of glycine betaine uptake, and this has been considered at odds with the fact that *E. coli*’s similar glycine betaine transport system uses a binding protein with a dissociation constant in the micromolar regime (Noell & Giovannoni, 2019). Our model shows that a nanomolar *K*_m_ can be achieved by a cell using a binding protein with a micromolar *K*_D_, if the cell has abundant binding proteins. We thus postulate that *P. ubique* relies on abundant binding proteins to achieve the high specific affinities that make it so successful in the vast nutrient-poor expanses of the ocean (Giovannoni, 2017). Although our model cannot rule out the alternative possibility that oligotrophs evolved binding proteins with lower *K*_D_ values to achieve very low *K*_M_ values, it does demonstrate that this is not required.

It has been previously hypothesized that oligotrophs achieve higher specific affinities than copiotrophs by having higher ratios of transport units to metabolic enzymes (Button, 1991), and this theory is still often used to explain the nutrient acquisition strategy of oligotrophs (Giovannoni, 2017). Here we propose an alternative theory: unlike cells using PTS, cells using ABC transport are able to increase their specific affinity without increasing their maximal uptake rate. As a result, oligotrophs may be able to closely match their transport and metabolic capacities. We hypothesize that it is for this reason that oligotrophs experience large nutrient upshifts as toxic (Schut *et al*, 1993; Koch, 2001; Cho & Giovannoni, 2004): since transport capacity rarely exceeds metabolic capacity, they may not have excretion systems as they would not be needed in the nutrient-poor ocean. Consequently, an atypical, large nutrient upshift would overwhelm the cytoplasm with substrate that the cell can neither process nor excrete.

Our metabolic model indicates that ABC transport systems are more efficient than PTS at low nutrient concentrations because expressing an additional binding protein has a lower proteomic cost than expressing an additional transport unit and, furthermore, does not incur real-estate costs on the inner membrane. Instead, the binding protein abundance is subject only to a constraint on the periplasmic density, a constraint that a cell can mitigate by modifying the fraction of its volume devoted to the periplasm. Our model predicts that the optimal periplasmic volume fraction increases as extracellular nutrient concentration decreases (Supplementary Figure S15): observations suggesting that the periplasm occupies up to 70% of the volume of an oligotrophic SAR11 *Pelagibacter* cell (Zhao *et al*, 2017) are in line with this prediction. Therefore, our model predicts that a majority of an oligotroph’s proteome is comprised of binding proteins (Figure 3*A*). This prediction is corroborated by metaproteomic analyses showing that binding proteins are among the most prevalent bacterial proteins found in the oligotrophic ocean (Sowell *et al*, 2009).

Taken together, our results provide a mechanistic explanation for the long-standing hypothesis of a rate–affinity trade-off for nutrient uptake by marine bacteria (Gudelj *et al*, 2007; Bosdriesz *et al*, 2018). The oligotroph’s reliance on binding proteins precludes its ability to attain high growth rates because of the slow diffusion of binding proteins in a large periplasm. Thus, a high substrate affinity comes at the cost of a reduced uptake rate. We find that the mechanism of this trade-off explains observations that the surface-area-to-volume ratio of a typical oligotroph is at least fivefold greater than that of a typical copiotroph (Ghuneim *et al*, 2018). We thus propose that the high translocation rates of PTS in copiotrophs are advantageous not only because they support greater uptake rates at high nutrient concentrations but also because these higher uptake rates support larger optimal cell volumes. This is of particular importance to motile copiotrophs, which must be large enough to overcome rotational diffusion in order to swim effectively toward nutrient hotspots (Mitchell, 1991). In addition, motile cells may not be able to attain values of *K*_M_ as low as those of oligotrophs because of the large periplasmic volumes that this requires. Because the distance between the outer and inner membranes dictates the length of the flagellar rotor, periplasmic volume is carefully regulated in motile cells (Miller & Salama, 2018) and typically does not exceed 20% of the cell volume (Oliver, 1996).

In summary, our work suggests that the constraints imposed by a rate–affinity trade-off between PTS and ABC transport systems shaped the divergent evolution of copiotrophic and oligotrophic bacteria in the ocean. By quantifying this trade-off, our model helps predict the achievable nutrient uptake rates and affinities of marine heterotrophic bacteria. These mechanistic predictions can be used to constrain the parametrizations of marine microbial ecosystem models used to understand how bacterial population dynamics may affect carbon flux rates in a changing ocean.

## Acknowledgements

This work was supported by a grant from the Simons Foundation (542395 to R.S. and 542389 to N.L.), as part of the Principles of Microbial Ecosystems (PriME) Collaborative, and by a Gordon and Betty Moore Foundation Investigator Award (GBMF3783 to R.S.). Contributions to the editing of this paper by Dr. Russell Naisbit are gratefully acknowledged. The authors thank Terry Hwa and Cameron Thrash for discussions and feedback.

## Author Contributions

NN, VF, and RS conceived the idea; NN and NL designed and developed model; NN performed optimizations, derived analytical results, and analyzed results; NN, NL, VF, and RS interpreted results; and NN, NL, and RS wrote the paper.

