## Supplemental Material for "Mechanistic model of nutrient uptake explains dichotomy between marine oligotrophic and copiotrophic bacteria"

Noele Norris, Naomi M. Levine, Vicente I. Fernandez, Roman Stocker

### Contents

**1 Modeling cytoplasmic transport** **2**
1.1 PTS transport . . . . . 2
1.2 ABC transport . . . . . 2

**2 Approximating ABC transport dynamics in Michaelis-Menten form** **4**

**3 *E. coli*'s ABC maltose transport system** **5**

**4 A simple cellular metabolic model** **9**

**5 Accounting for the energetic costs of transport** **14**

**6 References** **15**

**7 Additional figures** **19**

### 1 Modeling cytoplasmic transport

#### 1.1 PTS transport

We describe transport into the cell by a PTS by the following standard reactions and mass-action kinetic rates [2, 33]:

- $S_p + T \underset{k_{1r}}{\overset{k_{1f}}{\rightleftharpoons}} T{:}S$ :
  The periplasmic substrate, $S_p$, reversibly associates with the membrane-bound transport unit to form T:S.

- $T{:}S \xrightarrow{k_2} T + S_c$ :
  The substrate is actively transported into the cytoplasm.

The corresponding reaction rates are:

$$v_1 = k_{1f}[T][S]_p - k_{1r}[T{:}S], \tag{S-1}$$

$$v_2 = k_2[T{:}S]. \tag{S-2}$$

In addition is the conservation equation:

$$[T]_{total} = [T] + [T{:}S]. \tag{S-3}$$

At steady state, all of the concentrations remain constant so that $v_1 = v_2$. Thus, by Equations S-1, S-2, and S-3,

$$v_2 = [T]_{total} \frac{[S]_p}{\frac{k_2 + k_{1r}}{k_{1f}} + [S]_p} \tag{S-4}$$

To limit the number of free parameters, we take $k_2 = k_2 + k_{1r}$, which is equivalent to assuming that both reaction steps proceed irreversibly.

#### 1.2 ABC transport

We describe ABC transport by the following reactions and mass-action kinetic rates, which are similar to the description of transport via binding proteins in [12]:

- $S_p + BP \underset{k_{0r}}{\overset{k_{0f}}{\rightleftharpoons}} S{:}BP$ :
  The periplasmic substrate, $S_p$, reversibly associates with the binding protein, BP, to form ligand-binding protein complex, S:BP.

- $S{:}BP + T \underset{k'_{1r}}{\overset{k'_{1f}}{\rightleftharpoons}} T{:}S{:}BP$ :
  The substrate-binding protein complex reversibly associates with the membrane-bound ABC transport unit to form T:S:BP.

- $T{:}S{:}BP \xrightarrow{k'_2} T{:}BP + S_c$ :
  The substrate is actively transported into the cytoplasm leaving behind the binding protein in complex with the ABC transport unit, T:BP.

- $T{:}BP \underset{k_{3r}}{\overset{k_{3f}}{\rightleftharpoons}} T + BP$ :
  The binding protein reversibly associates with the ABC transport unit.

The corresponding reaction rates are:

$$v_1 = k_{0f}[\mathrm{S}]_\mathrm{p}[\mathrm{BP}] - k_{0r}[\mathrm{S{:}BP}], \tag{S-5}$$
$$v_2 = k'_{1f}[\mathrm{T}][\mathrm{S{:}BP}] - k'_{1r}[\mathrm{T{:}S{:}BP}], \tag{S-6}$$
$$v_3 = k'_2[\mathrm{T{:}S{:}BP}], \tag{S-7}$$
$$v_4 = k_{3f}[\mathrm{T{:}BP}] - k_{3r}[\mathrm{T}][\mathrm{BP}]. \tag{S-8}$$

At steady state, all of the concentrations remain constant:

$$0 = \frac{d[\mathrm{BP}]}{dt} = -v_1 + v_4, \tag{S-9}$$
$$0 = \frac{d[\mathrm{S{:}BP}]}{dt} = v_1 - v_2, \tag{S-10}$$
$$0 = \frac{d[\mathrm{T{:}S{:}BP}]}{dt} = v_2 - v_3, \tag{S-11}$$
$$0 = \frac{d[\mathrm{T{:}BP}]}{dt} = v_3 - v_4 \tag{S-12}$$
$$0 = \frac{d[\mathrm{T}]}{dt} = -v_2 + v_4. \tag{S-13}$$

Therefore, $v_1 = v_2 = v_3 = v_4$.

In addition are two conservation equations:

$$[\mathrm{BP}]_\mathrm{total} = [\mathrm{BP}] + [\mathrm{S{:}BP}] + [\mathrm{T{:}S{:}BP}] + [\mathrm{T{:}BP}] \text{ and} \tag{S-14}$$
$$[\mathrm{T}]_\mathrm{total} = [\mathrm{T}] + [\mathrm{T{:}S{:}BP}] + [\mathrm{T{:}BP}]. \tag{S-15}$$

We assume that the binding protein is more likely to bind to the transport units when bound to the substrate and that the unbound binding protein falls very quickly from the transport unit [17] so that $[\mathrm{T{:}BP}] \ll [\mathrm{T}]$. Therefore, because $v_2 = v_3$,

$$[\mathrm{T{:}S{:}BP}] \approx [\mathrm{T}]_\mathrm{total}\frac{[\mathrm{S{:}BP}]}{K'_\mathrm{T} + [\mathrm{S{:}BP}]}, \qquad K'_\mathrm{T} = \frac{k'_2 + k'_{1r}}{k'_{1f}}. \tag{S-16}$$

As we did for our model of PTS, we subsume $k'_{1r}$ into $k'_2$ to limit free parameters.

Because $v_1 = v_2$,

$$[\mathrm{S}]_\mathrm{p}[\mathrm{BP}] = K_\mathrm{D}[\mathrm{S{:}BP}] + \frac{k'_{1f}}{k_{0f}}[\mathrm{T}][\mathrm{S{:}BP}] - \frac{k'_{1r}}{k_{0f}}[\mathrm{T{:}S{:}BP}], \tag{S-17}$$

where $K_\mathrm{D} = k_{0r}/k_{0f}$. For effective translocation, $k'_{1f}[\mathrm{T}][\mathrm{S{:}BP}] > k'_{1r}[\mathrm{T{:}S{:}BP}]$. We assume that it is in fact much larger so that

$$[\mathrm{S}]_\mathrm{p}[\mathrm{BP}] \approx \left(K_\mathrm{D} + \frac{k'_{1f}}{k_{0f}}[\mathrm{T}]\right)[\mathrm{S{:}BP}]. \tag{S-18}$$

We therefore use the following system of equations to solve for the cytoplasmic uptake rate of an ABC transport system:

$$v_{\mathrm{c,\ ABC}} = k'_2[\mathrm{T}]_\mathrm{total}\frac{[\mathrm{S{:}BP}]}{K'_\mathrm{T} + [\mathrm{S{:}BP}]} \tag{S-19}$$
$$[\mathrm{BP}]_\mathrm{total} = [\mathrm{BP}] + [\mathrm{S{:}BP}] + [\mathrm{T{:}S{:}BP}] \tag{S-20}$$
$$[\mathrm{T}]_\mathrm{total} = [\mathrm{T}] + [\mathrm{T{:}S{:}BP}] \tag{S-21}$$
$$[\mathrm{S}]_\mathrm{p} = \left(K_\mathrm{D} + \frac{k'_1[\mathrm{T}]}{k_{0f}}\right)[\mathrm{S{:}BP}] \tag{S-22}$$

### 2 Approximating ABC transport dynamics in Michaelis-Menten form

In our model of ABC transport, the cytoplasmic uptake rate is the solution to the following set of equations:

$$v_{\mathrm{c,ABC}} = k_3[\mathrm{T{:}S{:}BP}] = k_3[\mathrm{T}]_{\mathrm{total}}\frac{[\mathrm{S{:}BP}]}{K_{\mathrm{T}} + [\mathrm{S{:}BP}]}, \quad K_{\mathrm{T}} = \frac{k_3}{k_2} \tag{S-23}$$

$$[\mathrm{BP}]_{\mathrm{total}} = [\mathrm{BP}] + [\mathrm{S{:}BP}] + [\mathrm{T{:}S{:}BP}], \tag{S-24}$$

$$[\mathrm{T}]_{\mathrm{total}} = [\mathrm{T}] + [\mathrm{T{:}S{:}BP}], \tag{S-25}$$

$$k_{1\mathrm{f}}[\mathrm{S}]_{\mathrm{p}}[\mathrm{BP}] = \left(k_{1\mathrm{r}} + k_2[\mathrm{T}]\right)[\mathrm{S{:}BP}]. \tag{S-26}$$

While we use the exact solution to these set of equations when running the proteome allocation problem, we can gain insight into how ABC transport achieves low affinities using an approximation that allows us to write the solution in Michaelis-Menten form.

We assume that $k_1[\mathrm{T}] \ll k_{0\mathrm{r}}$ (for our model, this implies $[\mathrm{T}] \ll 500\ \mu\mathrm{M}$, which holds for all of our solutions), so that we can simplify Equation S-26 to the following:

$$k_{0\mathrm{r}}[\mathrm{S{:}BP}] \approx k_{0\mathrm{f}}[\mathrm{S}]_{\mathrm{p}}\left([\mathrm{BP}]_{\mathrm{total}} - [\mathrm{S{:}BP}] - [\mathrm{T{:}S{:}BP}]\right).$$

Our optimal solutions and previous work on binding proteins in [12] suggest that the total abundance of binding proteins should exceed the total abundance of transport units for ABC transport to be effective; that is, $[\mathrm{BP}]_{\mathrm{total}} > [\mathrm{T}]_{\mathrm{total}}$. We additionally assume that $[\mathrm{BP}]_{\mathrm{total}} \gg [\mathrm{T{:}S{:}BP}]$, as observed, for example, in *E. coli*'s maltose transport system (see Section 3). Thus, we assume:

$$k_{0\mathrm{r}}[\mathrm{S{:}BP}] \approx k_{0\mathrm{f}}[\mathrm{S}]_{\mathrm{p}}\left([\mathrm{BP}]_{\mathrm{total}} - [\mathrm{S{:}BP}]\right).$$

Solving, we obtain:

$$[\mathrm{S{:}BP}] \approx [\mathrm{BP}]_{\mathrm{total}}\frac{[\mathrm{S}]_{\mathrm{p}}}{K_{\mathrm{BP}} + [\mathrm{S}]_{\mathrm{p}}}, \quad K_{\mathrm{D}} = \frac{k_{0\mathrm{r}}}{k_{0\mathrm{f}}}.$$

Plugging this solution for the periplasmic concentration of substrate bound to binding protein into Equation S-23, we obtain the Michaelis-Menten approximation:

$$v_{\mathrm{c,ABC}} \approx k_2'[\mathrm{T}]_{\mathrm{total}}\left(\frac{[\mathrm{BP}]_{\mathrm{total}}}{K_{\mathrm{T}}' + [\mathrm{BP}]_{\mathrm{total}}}\right)\left(\frac{[\mathrm{S}]_{\mathrm{p}}}{\frac{K_{\mathrm{T}}'K_{\mathrm{D}}}{K_{\mathrm{T}}' + [\mathrm{BP}]_{\mathrm{total}}} + [\mathrm{S}]_{\mathrm{p}}}\right). \tag{S-27}$$

In Figure S-6, we compare this approximation with the exact solutions to the system of four equations for a variety of parameter values.

### 3 *E. coli*'s ABC maltose transport system

Because the maltose transport system of *E. coli* is among the best-studied ABC transport systems, we use it to validate our simple model of ABC transport. Previous work concluded that an ABC transport model in which $K_{\mathrm{M}}$ could become substantially lower than $K_{\mathrm{D}}$ was invalid because it did not capture the behavior of *E. coli*'s maltose transport system [9]. However, here we show that it can in fact describe the behavior once we account for the effects of maltoporin limitations on transport in the micromolar regime, as has been observed experimentally [14, 53]. Thus, although *E. coli*'s maltose transport system reaches a $K_{\mathrm{M}}$ only twofold lower than the $K_{\mathrm{D}}$ of its binding protein, we find that this is because the cell's affinity is limited by the permeability of maltose through the cell's outer membrane.

We use the equations derived in Section 1 (Equations S-19 to S-22) to model the cytoplasmic transport rate of maltose from the periplasm ($\mathrm{S_p}$) into the cytoplasm. We model the periplasmic transport rate ($\mathrm{S_p} \rightarrow \mathrm{S_{ext}}$) as the summation of the transport rates due to the maltoporin, LamB, and the general porin, OmpF [3]. While the diffusion of maltose into the periplasm via a general porin can be described by a simple diffusive uptake term that is proportional to the difference of the extracellular maltose concentration and the periplasmic concentration of free maltose, transport by facilitated diffusion by specific porins is better modeled using a half-saturation concentration [13]. Therefore, we model the periplasmic uptake rate as:

$$v_p = V_{\mathrm{LamB}} \frac{[\mathrm{S}]_{\mathrm{ext}} - [\mathrm{S}]_{\mathrm{p}}}{K_{\mathrm{LamB}} + [\mathrm{S}]_{\mathrm{ext}} + [\mathrm{S}]_{\mathrm{p}}} + V_{\mathrm{OmpF}} \left([\mathrm{S}]_{\mathrm{ext}} - [\mathrm{S}]_{\mathrm{p}}\right) \tag{S-28}$$

To solve for the effective rate of transport of maltose at a specified concentration of extracellular maltose, we solve for the value of $[\mathrm{S}]_{\mathrm{p}}$ such that $v_p = v_c$ and plug this value into $v_c$ to solve for the total uptake rate. We vary the concentrations of extracellular maltose and use MATLAB's function *fit* to determine the Michaelis-Menten kinetic values, $\mathrm{V_{max}}$ and $\mathrm{K_M}$, that best fit our solution for the total uptake rate:

$$v_c = V_{\mathrm{max}} \frac{[\mathrm{S}]_{\mathrm{ext}}}{K_{\mathrm{M}} + [\mathrm{S}]_{\mathrm{ext}}}. \tag{S-29}$$

For all parameter values used in our model, we use estimates from previous experimental work:

- $[\mathrm{BP}]_{\mathrm{total}} = 1$ mM. Estimate from [10] and references therein.
- $[T]_{\mathrm{total}} = 10\ \mu$M. We assume that there are 100 times more binding proteins than transport units [46].
- $K_{\mathrm{D}} = 2\ \mu$M. Estimate from [10] and references therein.
  - $k_{0\mathrm{f}} = 10^5$ mM$^{-1}$ sec$^{-1}$. Estimate from [37].
  - $k_{0\mathrm{r}} = 200$ sec$^{-1}$. (Estimate from [37]. $K_{\mathrm{D}} = k_{0\mathrm{r}}/k_{0\mathrm{f}}$)
- $K_{\mathrm{T}} = 100\ \mu$M. Estimate from [5].
  - $k_2' = 10$ sec$^{-1}$. Estimate from [30] and references therein.
  - $k_1' = 0.1$ mM$^{-1}$sec$^{-1}$. ($K_{\mathrm{T}} = k_2'/k_1'$)
- $K_{\mathrm{LamB}} = 10$ mM. Estimate from [3].

- $V_{\text{LamB}} = 10$ mM/sec. We use the estimate that $V_c/V_{\text{LamB}} = 10^{-4}$, where the maximal cytoplasmic uptake rate $V_c = k_2'[\text{T}]_{\text{total}}$, so that transport becomes porin-limited at approximately 1 $\mu$M, as shown in figure below.

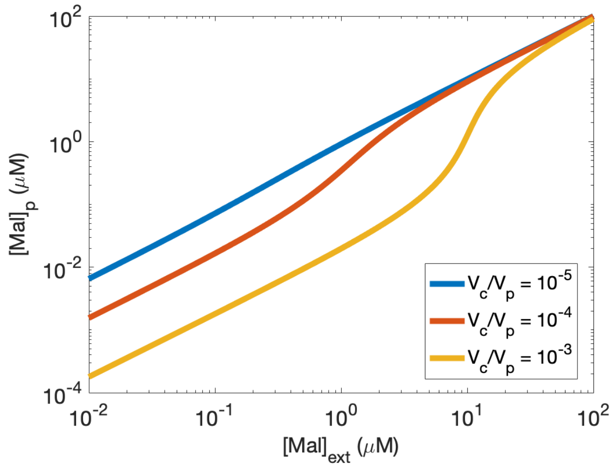

- $V_{\text{OmpF}} = 0.1$ sec$^{-1}$. We estimate that $V_{\text{OmpF}} = 0.1 \times V_{\text{LamB}}/K_{\text{LamB}}$, as observed in [3].

To obtain the plots shown below, we varied the abundances of either maltoporins or binding proteins to match previous experiments.

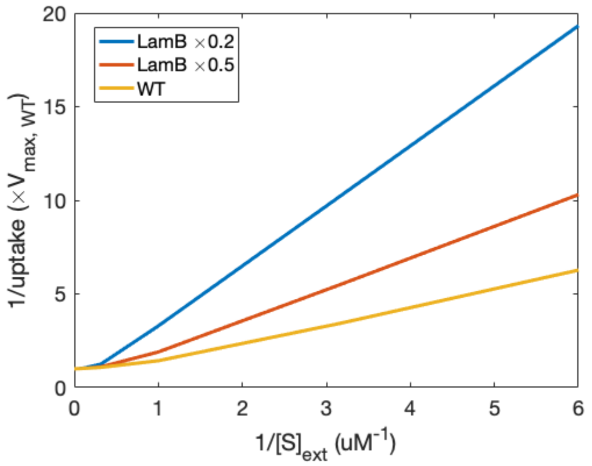

Figure S-1: **Effects of maltoporin abundance on uptake.** To replicate Fig. 6 in [14], we solve for the total uptake rate assuming wild-type maltoporin levels or twofold or fivefold reductions in maltoporin levels. We assume that $V_{\text{LamB}}$ scales proportionally with maltoporin abundance. Uptake rates are scaled by the maximal uptake rate of the wild type, $V_{\text{max,WT}}$, so that the lines cross the y-axis at 1.

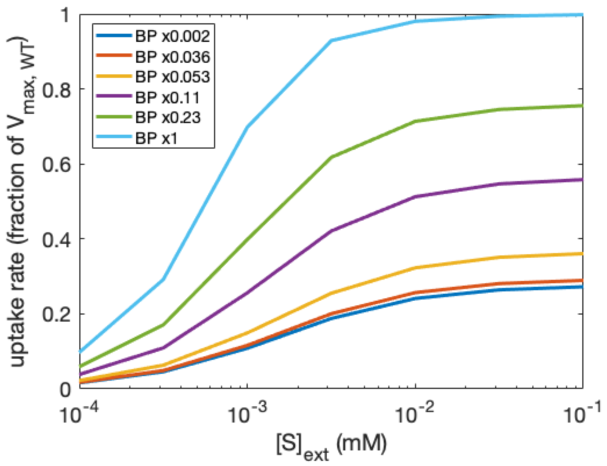

Figure S-2: **Effects of binding protein abundance on uptake.** To replicate Fig. 4 in [35], we solve for the total uptake rate assuming various fractions of the wild-type MalE abundances, corresponding to the abundances estimated in the various mutants used in [35]. Our model replicates the sigmoidal dependence of uptake on maltose concentration.

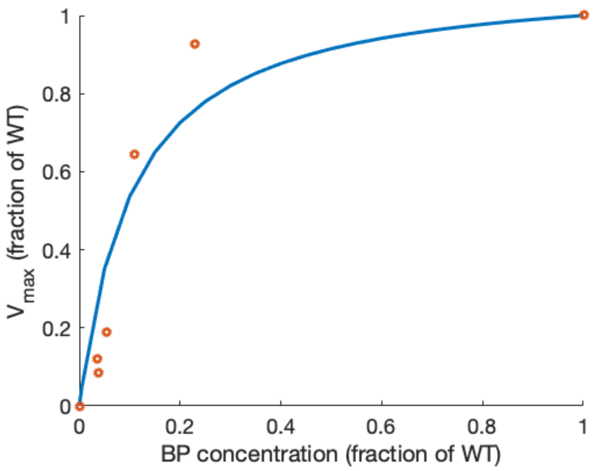

Figure S-3: **Effects of binding protein abundance on maximal uptake rate.** To replicate Fig. 6A in [35], we calculate the estimated uptake rates for various MalE abundances at an extracellular maltose concentration of 100 mM and take this to be the maximal uptake rate. To contrast our estimates with the values obtained in [35], we scale both our model estimates and their data so that the maximal uptake rate of the wild-type is 1. Our model well captures all the data points except for the response with a binding protein abundance of 0.23× wild-type abundance. It is unclear what causes this discrepancy. Our model assumptions may be incorrect and thus cannot capture the steep saturation in response. However, the discrepancy may also be due to the fact that [35] does not compare measurements against a wild-type strain but only uses strains that constitutively express the global activator of the *mal* regulon, MalT. Thus, there are likely disparities between the Mal protein abundances of the cells used in their experiments and the measured wild-type abundances we use in our model estimates.

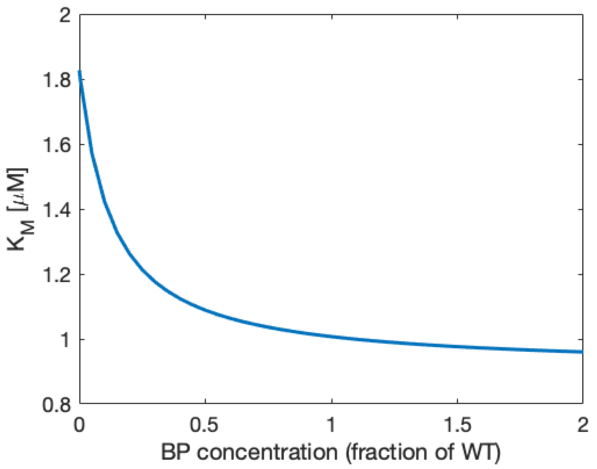

Figure S-4: **Effects of binding protein abundance on the half-saturation concentration of uptake.** Our model does a particularly good job of capturing the dependence of the effective half-saturation concentration of maltose uptake on binding protein abundance, as observed in [35]. Both our model estimates and [35] show that the effective $K_{\mathrm{M}}$ value of total maltose uptake asymptotes at a value of approximately 1 $\mu$M as MalE abundance increases.

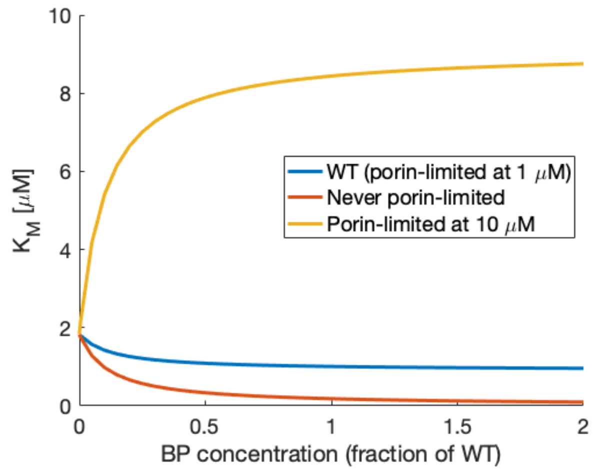

Figure S-5: **The permeability of the outer membrane limits half-saturation concentration of uptake.** Previous modeling work [9] assumed that the dissociation constant of MalE to maltose was $K_{\mathrm{D}} = 1$ $\mu$M (whereas more recent measurements suggest a $K_{\mathrm{D}}$ value of 2$\mu$M [37]) and concluded from the observations in [35] that the effective $K_{\mathrm{M}}$ of an ABC transport system could never be lower than the dissociation constant $K_{\mathrm{D}}$ of the system's binding protein to its substrate. However, the work erroneously assumed that maltose uptake is never porin-limited and thus conflated the effective half-saturation concentration of total maltose uptake with the half-saturation concentration of cytoplasmic uptake. Our model instead suggests that the outer membrane permeability limits uptake in the micromolar regime so that the the half-saturation concentration of uptake is a function both of binding protein abundance and maltoporin abundance. Indeed, if maltose transport is never porin-limited, our model suggests that the effective half-saturation concentration could be almost a factor of ten lower than the wild-type.

### 4 A simple cellular metabolic model

#### 4.1 Proteome allocation problem

To compare the performance of PTS and ABC transport, we incorporate models of them into a single-cell metabolic model that is an extension of the self-replicator model proposed by Molenaar and others [39]. We use this model to solve a proteome allocation problem that maximizes the steady-state, exponential growth rate in a homogeneous environment with a specified external concentration of a generic carbon substrate that we assume provides the cell with everything it needs to replicate. We optimize over the following cell characteristics: surface-area-to-volume ratio, periplasmic volume fraction, intracellular metabolite concentrations, and fractions of the proteome devoted to particular protein groups. Below we fully specify the problem. The values in blue are the independent variables that our problem optimizes over. The values in red are functions of the independent variables. Definitions of each variable are given below.

$$\underset{x}{\text{maximize}}\ \boldsymbol{\mu} \quad \text{subject to: Eq.C 1-7, Ineq.C 1-3, and } x_i \geq 0, \forall i,$$

$$\text{where } x = (x_{\text{m}}, \phi, r, f_{\text{p}}, \boldsymbol{\mu}).$$

$$x_{\text{m}} = [[\text{S}]_{\text{p}}, [\text{S}]_{\text{c}}, [\text{A}], [\text{W}], [\text{P}]]$$

$$\phi = [\phi_{\text{BP}}, \phi_{\text{T}}, \phi_{\text{E}}, \phi_{\text{W}}, \phi_{\text{R}}]$$

Eq.C 1-5 (Steady-state growth) :
$$\frac{dx_{\text{m}}}{dt} = N v_{\text{r}} - \boldsymbol{\mu} x_{\text{m}} = 0$$

$$v_r = \begin{bmatrix} v_{\text{diff}} & (\text{S}_{\text{ext}} \to \text{S}_{\text{p}}) \\ v_{\text{c}} & (\text{S}_{\text{p}} \to \text{S}_{\text{c}}) \\ k_{\text{E}}[\text{E}]\frac{[\text{S}]_{\text{c}}}{K_{\text{M,E}}+[\text{S}]_{\text{c}}} & (5\text{S}_{\text{c}} \to 6\text{A}) \\ k_{\text{W}}[\text{M}]\left(\frac{[\text{A}]}{K_{\text{M,W}}+[\text{A}]}\right) & (\text{A} \to \text{W}) \\ k_{\text{R}}[\text{R}]\left(\frac{[\text{A}]}{K_{\text{M,R}}+[\text{A}]}\right) & (\text{A} \to \text{P}) \end{bmatrix},$$

where the components of the proteome($\mathcal{P}$) are: $[\text{P}]_j = \phi_j \alpha_j [\text{P}]$.

Eq.C 6 (Proteome allocation) :

$$1 = \sum_{i \in \mathcal{P}} \phi_i$$

Eq. C 7 (Cell membranes) :

$$4\pi \left(1 + (1 - f_{\text{p}})^{2/3}\right) r^2 = [\text{W}] \left(4/3\pi r^3\right) a_{\text{W}}$$

Ineq. C 1 (Cytoplasmic cell density) :

$$\sum_{j \in \mathcal{M}_{cyto}} m_j x_{\text{m}}(j) \leq \rho$$

Ineq. C 2 (Periplasmic cell density) :

$$\sum_{j \in \mathcal{M}_{peri}} m_j x_{\text{m}}(j) \leq \rho$$

Ineq. C 3 (Inner Membrane "Real Estate") :

$$f_{\text{SA}} \left(4\pi(1 - f_{\text{p}})^{2/3} r^2\right) \geq [\text{T}] \left(4/3\pi r^3\right) a_{\text{T}}$$

Note that all components of the proteome, $[\mathrm{P}]_i$, are in units of abundance divided by cytoplasmic volume. The periplasmic concentration of the binding proteins is thus, $(1 - f_\mathrm{p})[\mathrm{BP}]/f_\mathrm{p}$, where $f_\mathrm{p}$ is the fraction of the cell volume comprised of the periplasm.

This proteome allocation problem optimizes over the following variables to determine the maximal exponential growth rate, $\mu$, given a particular extracellular carbon substrate concentration, $[\mathrm{S}]_\mathrm{ext}$:

- the intracellular metabolite concentrations, $x_\mathrm{m}$, which include:
  - $[\mathrm{S}]_\mathrm{p}$, the periplasmic concentration of generic carbon substrate;
  - $[\mathrm{S}]_\mathrm{c}$, the cytoplasmic concentration of generic carbon substrate;
  - [A], the cytoplasmic concentration of the precursor building block of both proteins and the cell membranes, presumed to be a generic amino acid;
  - [W], the number of generic cell membrane units divided by the cytoplasmic volume of the cell;
  - [P], the number of amino acids incorporated into proteins divided by the cytoplasmic volume of the cell;
- the proteome fractions, $\phi$, which include:
  - $\phi_\mathrm{BP}$, the fraction of the proteome comprised of periplasmic binding proteins that enable ABC transport (for PTS transport, $\phi_\mathrm{BP} = 0$);
  - $\phi_\mathrm{T}$, the fraction of the proteome comprised of inner-membrane-bound transport units that translocate the periplasmic carbon substrate, $\mathrm{S_p}$ into the cytoplasm ($\mathrm{S_c}$);
  - $\phi_\mathrm{E}$, the fraction of the proteome comprised of metabolic enzymes that transform the cytoplasmic substrate, $\mathrm{S_c}$, into a generic amino acid building block, A;
  - $\phi_\mathrm{W}$, the fraction of the proteome comprised of membrane biosynthesis enzymes that build both the inner and outer membranes, W, from the building block A;
  - $\phi_\mathrm{R}$, the fraction of the proteome comprised of ribosomes that use building block A to elongate peptide chains and thus express the proteome, P;
- the cell radius, $r$, assuming a spherical cell; and
- the periplasmic volume fraction, $f_\mathrm{p}$.

To solve for these optimal parameter values, the optimization problem is subject to a variety of constraints:

- The equality constraints 1-5 are ordinary differential equations that assume a balanced, steady-state exponential growth of each of the cellular components. $N$ is thus a stoichiometry matrix that describes how many of each input metabolite is needed to form an output, and $v_\mathrm{r}$ is a vector of Michaelis-Menten reaction rates. Explanations for each parameter choice for the stoichiometry matrix and reaction rates are provided below.
- Equality constraint 6 specifies that the proteome fractions must sum to one.
- Equality constraint 7 specifies that the amount of cell membrane units expressed must cover both the inner and outer membranes of the cell.
- Inequality constraint 1 specifies that the sums of the molecular weights of each cytoplasmic metabolite ($\mathrm{S_c}$, A, $(\phi_\mathrm{E} + \phi_\mathrm{W} + \phi_\mathrm{R})$P) multiplied by its corresponding cytoplasmic concentration must be less than or equal to the maximal allowed density, $\rho$.

- Inequality constraint 2 specifies that the sums of the molecular weights of each periplasmic component ($S_p$, BP) multiplied by its corresponding periplasmic concentration must be less than or equal to the maximal allowed density, $\rho$.
- Inequality constraint 3 specifies that the inner-membrane bound transport units, T, must fit on the surface area of the inner membrane.

For reasonable parameter values, we use glucose consumption by *E. coli* as a model uptake system.

#### 4.2 Periplasmic uptake

$$\mathrm{S_{ext}} \xrightarrow{v_\mathrm{p}} \mathrm{S_p}$$

The rate of uptake of the carbon substrate from the extracellular environment with concentration $[\mathrm{S}]_\mathrm{ext}$ into the periplasm is a function of both the rate of the diffusion of the substrate to the cell's surface and the rate of the diffusion of the substrate through porins on the outer membrane from the surface directly into the periplasm. To reduce the complexity of the model and focus on the implications of variations of cytoplasmic transport, we assume in this work that the periplasmic uptake rate is set simply by the rate of the diffusion of the substrate to the cell's surface and not by the abundance of expressed porins on the outer membrane. (In [43], we expanded the model to account for porins and this did not change any of our qualitative results.)

Thus, we assume that [4]:

$$v_\mathrm{diff} = 3D\frac{[\mathrm{S}]_\mathrm{ext} - [\mathrm{S}]_\mathrm{p}}{f_\mathrm{p} r^2}, \qquad \left[\frac{\text{Periplasmic Concentration}}{\text{Time}}\right]$$

where $[\mathrm{S}]_\mathrm{ext}$ is the concentration of the substrate in the external environment; $[\mathrm{S}]_\mathrm{p}$ is the concentration of substrate in the periplasm; $r$ is the cell's radius; $D$ is the diffusivity of the substrate in the external environment; and $f_\mathrm{p}$ is the fraction of the volume of the cell that is comprised of the periplasm.

In our baseline model, we assume that $D = 10^{-8}$ cm$^2$ msec$^{-1}$, which is approximately the diffusivity of a glucose molecule in water.

We run the optimization problem over a range of external concentrations, from $[\mathrm{S}]_\mathrm{ext} = 10^{-6}$ mM to $[\mathrm{S}]_\mathrm{ext} = 100$ mM.

#### 4.3 Cytoplasmic uptake

$$\mathrm{S_p} \xrightarrow{v_\mathrm{c}} \mathrm{S_c}$$

We run the optimization problem using one of two models for cytoplasmic uptake. Both models are explained and derived within the manuscript.

##### 4.3.1 PTS transport

For PTS transport, we use the following equation:

$$v_\mathrm{c,\ PTS} = k_2[\mathrm{T}]\frac{[\mathrm{S}]_\mathrm{p}}{K_\mathrm{T} + [\mathrm{S}]_\mathrm{p}},$$

where [T] is the number of membrane-bound transport units divided by the cytoplasmic volume of the cell, $k_2$ is the translocation rate of the transport-unit, and $K_\mathrm{T}(= k_2/k_1)$ is the half-saturation constant of the transport unit.

In our baseline model, we use:

- $k_2 = 200\ \mathrm{sec}^{-1}$. The translocation rate of the glucose transport ptsI is approximately 210 molecules/sec. (BNID:103693, [8])
- $K_\mathrm{T} = 10\ \mu\mathrm{M}$. The effective half-saturation constant for glucose transport was observed to be between $5 - 20\ \mu\mathrm{M}$ [24].
- $\alpha_T = 1/1600$. The number of amino acids of each protein comprising the system are: 831 for PtsI, 85 for PtsH, 477 for PtsG, and 169 for Crr ([27], eco:02060).

##### 4.3.2 ABC transport

For ABC transport, we use the following equation, which is the solution to Eqs. 2-5 in the main text:

$$v_\mathrm{c,ABC} = \frac{k_2'}{2(k_2' + k_\mathrm{0f}[\mathrm{S}]_\mathrm{p})}\Big( \left[k_\mathrm{0r}K_\mathrm{T}' + k_\mathrm{0f}[\mathrm{S}]_\mathrm{p}\left([\mathrm{BP}]_\mathrm{peri} + [\mathrm{T}]_\mathrm{peri} + K_\mathrm{T}'\right) + k_2'[\mathrm{T}]_\mathrm{peri}\right] \ldots$$
$$- \sqrt{4[\mathrm{BP}]_\mathrm{peri}k_\mathrm{0f}K_\mathrm{T}'[\mathrm{S}]_\mathrm{p}(k_\mathrm{0r} + k_\mathrm{0f}[\mathrm{S}]_\mathrm{p}) + (k_\mathrm{0r}K_\mathrm{T}' + k_2'[\mathrm{T}]_\mathrm{peri} + k_\mathrm{0f}[\mathrm{S}]_\mathrm{p}(K_\mathrm{T}' + [\mathrm{T}]_\mathrm{peri} - [\mathrm{BP}]_\mathrm{peri}))^2}\ \Big),$$

where $[\mathrm{BP}]_\mathrm{peri}$ is the concentration of binding proteins in the periplasm; $[\mathrm{T}]_\mathrm{peri}$ is the number of membrane-bound transport units divided by the volume of the periplasm; and the transport rates and half-saturation constants are defined and parametrized below.

In our baseline model, we use the following values:

- The binding-protein dissociation constant $K_\mathrm{D} = 1\ \mu\mathrm{M}$. $K_\mathrm{D} = k_\mathrm{0r}/k_\mathrm{0f}$, where $k_\mathrm{0f} = 1 \times 10^5\ \mathrm{mM}^{-1}\mathrm{sec}^{-1}$ and $k_\mathrm{0r} = 100\ \mathrm{sec}^{-1}$. The *E. coli* maltose binding protein, MalE, has a dissociation constant of approximately $1\ \mu\mathrm{M}$ [23]. The association and dissociation rates, $k_\mathrm{0f}$ and $k_\mathrm{0r}$ respectively, were estimated from [37].
- The translocation rate $k_2' = 2\ \mathrm{sec}^{-1}$. As explained in the manuscript, the diffusivity of the binding protein-substrate complex is approximated to be one hundred times slower than the diffusivity of the substrate itself. Thus, we approximate that the transport rate for ABC transport is one hundred times lower than that of direct transport.
- The transport dissociation constant $K_T' = 10\mu M$. We assume that the transport unit dissociation constants for direct and ABC transport are the same. This implies that the binding-protein-substrate complex and transport unit association rate $k_1' = 200\ \mathrm{mM}^{-1}\mathrm{sec}^{-1}$.
- $\alpha_{BP} = 1/400$. MalE is comprised of 396 amino acids.
- $\alpha_T$ and $a_T$ are the same as for direct transport.

#### 4.4 Metabolism

$$5\mathrm{S_c} \xrightarrow{v_\mathrm{E}} 6\mathrm{A}$$

The set of metabolic enzymes, E, transform the intracellular nutrient, $\mathrm{S_c}$, into amino acids at a rate of

$$v_\mathrm{E} = k_\mathrm{E}[\mathrm{E}]\frac{[\mathrm{S}]_\mathrm{c}}{K_\mathrm{M,E} + [\mathrm{S}]_\mathrm{c}}.$$

For simplicity, we consider proteins to be made of just one type of amino acid, and use the following parameter values.

- The metabolic turnover rate is $k_\mathrm{E} = 100$ sec$^{-1}$. The phosphofructokinase reaction is a rate-limiting step of glycolysis ([33]) and has a turnover rate on the order of approximately 100 sec$^{-1}$ (EC 2.7.1.11, [15]).
- The half-saturation constant of metabolism $K_\mathrm{E} = 100$ $\mu$M. This is the approximate half-saturation constant of phosphofructokinase-2 for fructose 6-phosphate [15].
- $\alpha_E = 1/40{,}000$. There are about 130 genes in glycolysis and the biosynthesis of amino acids ([27], M00002, eco01230).

Because the different amino acids are composed of various amounts of carbon, we average over all of the amino acids and assume that 5 glucose molecules provide enough carbon to produce 6 amino acids.

#### 4.5 Membrane biosynthesis

$$\mathrm{A} \xrightarrow{v_\mathrm{M}} \mathrm{W}$$

We assume that the set of membrane biosynthesis enzymes, M, transform the amino acid building block into the components of both the inner and outer membrane of the cell at a rate of

$$v_\mathrm{M} = k_\mathrm{M}[\mathrm{M}]\frac{[\mathrm{A}]}{K_\mathrm{M} + [\mathrm{A}]}.$$

In our model, we use the following baseline parameter values:

- The membrane biosynthesis turnover rate $k_\mathrm{M} = 3$ sec$^{-1}$. This is the catalytic rate of penicillin-binding protein 1 [50].
- The half-saturation constant of membrane biosynthesis $K_\mathrm{M} = 20$ $\mu$M. This is the approximate half-saturation constant of MurC for L-alanine [52].
- $\alpha_\mathrm{M} = 1/21{,}000$. There are 19 enzymes involved in peptidoglycan biosynthesis ([27], eco:01011).

#### 4.6 Protein expression

$$\mathrm{A} \xrightarrow{v_\mathrm{R}} \mathrm{P}$$

The ribosomes, R, build the proteome, P, from the intracellular pool of the building block amino acid at a rate of

$$v_\mathrm{R} = k_\mathrm{R}[\mathrm{R}]\frac{[\mathrm{A}]}{K_\mathrm{R} + [\mathrm{A}]}$$

Note that we take as our unit of proteome a single amino acid so that [P] is the concentration of amino acids that comprise the proteome. We use the following baseline parameter values in our model:

- The elongation rate $k_\mathrm{R} = 15$ sec$^{-1}$. The peptide chain elongation rate is from about 12 to 20 amino acids per second per ribosome ([8], BNID:107868).

- The half-saturation constant of elongation $K_{\mathrm{R}} = 300$ $\mu$M. Estimates within a range of $200 - 400$ $\mu$M have been made for the half-saturation constant of several aminoacyl-tRNA synthetases to various amino acids [55].
- $\alpha_{\mathrm{R}} = 1/20{,}000$. A ribosome is comprised of both ribosomal protein (approximately 7,000 amino acids) and rRNA (approximately 5,000 ribonucleotides) [32].

##### 4.7 Density and surface area constraints

The abundances of the intracellular components are constrained by either the maximal allowed density of the cell or, for the membrane-bound components, the surface area of the cell. For Inequality Constraints 1-3, we use the following baseline parameters:

- $\rho = 0.3$ g/mL. The density of a cell is approximately 0.34 g/mL ([8], BNID:109049).
- $m_{\mathrm{A}} = 110$ Da. The average molecular weight of an amino acid is 110 Da.
- $m_{\mathrm{Sc}} = 180$ Da. The molecular weight of glucose is 180 Da.
- $m_{\mathrm{W}} = 220$ Da. The molecular weight of N-acetylglucosamine is 221 Da.
- $a_{\mathrm{T}} = 50$ nm$^2$. The surface area of one transport unit equals approximately 50nm$^2$ [38].
- $a_{\mathrm{W}} = 2$ nm$^2$. The surface area of one unit of peptidoglycan [54] .

For the baseline model, we assume that half of the surface areas of both the inner and outer membranes are available for proteins, $f_{\mathrm{SA}} = 0.5$.

##### 4.8 Solving the Optimization Problem

To solve the proteome allocation problem, we use MATLAB's constrained nonlinear multivariable function solver, **fmincon**. To ensure that our solutions are global and not simply local ones, we run the solver 50 times for each optimization problem, each time using different initial conditions. We additionally transform the units of both the constraints and variables so that their magnitudes are all approximately 1.

#### 5 Accounting for the energetic costs of transport

While this work focuses on the proteomic costs of particular traits, many traits also have an energetic cost. Cells maintain an ATP/ADP ratio about $10^8$ times greater than its equilibrium ratio to drive energetically unfavorable processes by coupling them with ATP hydrolysis [22, 31]. Thus, as the cell converts ATP to ADP to carry out energetically costly reactions, it must maintain energy homeostasis by continually using energy to convert ADP back into ATP. Thus, carbon is a crucial substrate for heterotrophic bacteria because it is used not only as a building block for amino acids and other biomass but is also a means for synthesizing ATP. Aerobic heterotrophs can use carbon to produce ATP via two main processes. First, the cells can perform glycolysis, a substrate-level phosphorylation within the cytoplasm in which a glucose molecule is transformed into pyruvate and generates two ATP molecules from ADP. In addition, by a second process, called oxidative phosphorylation or cellular respiration, the cell uses oxygen as an electron acceptor to further break down the pyruvate into carbon dioxide to create at most a total of 36 ATP molecules per molecule of glucose [44]. Cellular respiration is carried out by a cascade of proteins, called the electron transport chain, and ATP synthase. The electron transport chain units are bound to the inner membrane, as they generate an electrochemical proton gradient across the membrane, known as the proton-motive force. This force, via a translocation of protons, drives ATP synthase to produce ATP from ADP. Thus, the proton-motive force is an additional currency of energy used by the cell [22], and is, in fact, used directly as energy by a number of symporters.

In [43], we used an expansion of the metabolic model that additionally considered energetic cost. We specified the energetic cost of each protein group by quantifying the equivalent number of ATP used per second. The ABC transport system uses ATP directly. However, it is not known how tightly linked substrate translocation is to ATP usage [48]. While some studies suggest a ratio of 1-2 ATP units used per substrate translocated [40, 45], others suggest that ABC transport may be more energetically costly and that it may even use ATP in a futile cycle when no substrate is present [47]. On the other hand, it has been argued that PTS transport is the most energy efficient method of transport for carbohydrates because its energy-consuming phosphorylation step, in which a phosphoryl group from phosphoenolpyruvate is transferred to the sugar substrate, is also a required step in glycolysis [49]. The energetic cost of this phosphorylation is equivalent to one ATP consumed per substrate because the transformation of the phosphoenolpyruvate into pyruvate produces one ATP molecule [33].

This analysis suggests that ABC transport has both higher proteomic and energetic costs per substrate translocated than PTS transport systems. However, our analysis in [43] showed that the energetic cost did not affect our qualitative results (Fig. S-12). Therefore, here we simplified the model to only consider proteomic cost.

[6] Berg, Howard C. *Random Walks in Biology.* Princeton University Press, 1993.

[7] Berg, Jeremy M., John L Tymoczko, and Lubert Stryer. Biochemistry. 5th ed., W H Freeman, 2002.

[15] BRENDA: The Comprehensive Enzyme Information System. https://www.brenda-enzymes.org/

[16] G. M. Cooper, “Structure of the Plasma Membrane,” in *The Cell: A Molecular Approach. 2nd edition.* 2000. [Online]. Available: https://www.ncbi.nlm.nih.gov/books/NBK9898/ [Accessed: 21-June-2017]

[17] Davidson, A. L., Dassa, E., Orelle, C. & Chen, J. “Structure, Function, and Evolution of Bacterial ATP-Binding Cassette Systems.” *Microbiol. Mol. Biol. Rev.* 72, 317?364 (2008).

[18] Dusenbery, David B. *Living at Micro Scale: The Unexpected Physics of Being Small.* Harvard University Press, 2009.

[19] Erickson, David W., Severin J. Schink, Vadim Patsalo, James R. Williamson, Ulrich Gerland, and Terence Hwa. “A Global Resource Allocation Strategy Governs Growth Transition Kinetics of Escherichia Coli.” *Nature* 551, no. 7678 (November 2, 2017): 119-23.

[20] Etzold, C., et al. “Turnover Number of Escherichia Coli F0F1 ATP Synthase for ATP Synthesis in Membrane Vesicles.” European Journal of Biochemistry, vol. 243, no. 1?2, Jan. 1997, pp. 336-43.

[21] S. J. Giovannoni et al., “Genome Streamlining in a Cosmopolitan Oceanic Bacterium,” *Science*, vol. 309, no. 5738, pp. 12421245, Aug. 2005.

[22] Harold, F. M. “The Vital Force: A Study of Bioenergetics.” (W. H. Freeman, 1986).

[23] Hengge, Regine, and Winfried Boos. “Maltose and Lactose Transport in Escherichia Coli: Examples of Two Different Types of Concentrative Transport Systems.” Biochimica et Biophysica Acta, vol. 737, no. 3, Aug. 1983, pp. 443?78.

[24] K. Jahreis, E. F. Pimentel-Schmitt, R. Brckner, and F. Titgemeyer, “Ins and outs of glucose transport systems in eubacteria,” FEMS Microbiology Reviews, vol. 32, no. 6, pp. 891907, Nov. 2008.

[25] S. Jo, E. L. Wu, D. Stuhlsatz, J. B. Klauda, G. Widmalm, and W. Im, “Lipopolysaccharide Membrane Building and Simulation,” *Methods Mol. Biol.*, vol. 1273, pp. 391406, 2015.

[26] Kaleta, Christoph, Sascha Schuble, Ursula Rinas, and Stefan Schuster. “Metabolic Costs of Amino Acid and Protein Production in Escherichia Coli.” *Biotechnology Journal* 8, no. 9 (September 1, 2013): 110514.

[27] KEGG Genome Database. https://www.genome.jp/kegg/genome/html.

[33] A. L. Lehninger, D. L. Nelson, and M. M. Cox, Lehninger *Principles of Biochemistry.* Sixth Edition. Macmillan, 2012.

[34] W. Liebermeister, E. Noor, A. Flamholz, D. Davidi, J. Bernhardt, and R. Milo, “Visual account of protein investment in cellular functions,” *Proceedings of the National Academy of Sciences*, vol. 111, no. 23, pp. 84888493, Jun. 2014.

[35] Manson, M. D., Boos, W., Bassford, P. J. & Rasmussen, B. A. “Dependence of maltose transport and chemotaxis on the amount of maltose-binding protein.” J. Biol. Chem. 260, 9727?9733 (1985).

[36] A. G. Marr, “Growth rate of Escherichia coli.,” Microbiol Rev, vol. 55, no. 2, pp. 316333, Jun. 1991.

[37] Miller, D. M., et al. “Rates of Ligand Binding to Periplasmic Proteins Involved in Bacterial Transport and Chemotaxis.” Journal of Biological Chemistry, vol. 258, no. 22, Nov. 1983, pp. 13665?72.

[38] Milo, Ron, and Rob Phillips. *Cell Biology by the Numbers.* Garland Science, 2015.

[39] D. Molenaar, R. van Berlo, D. de Ridder, and B. Teusink, “Shifts in growth strategies reflect tradeoffs in cellular economics,” *Molecular Systems Biology*, vol. 5, no. 1, p. 323, Jan. 2009.

[40] Muir, M., Williams, L. & Ferenci, T. “Influence of Transport Energization on the Growth Yield of *Escherichia coli*.” J. BACTERIOL. 163, 6 (1985).

[41] Neidhardt, Frederick Carl, et al. Physiology of the Bacterial Cell: A Molecular Approach. Sinauer Associates, 1990.

[42] Nikaido, H. “Effect on Solute Size on Diffusion Rates through the Transmembrane Pores of the Outer Membrane of Escherichia Coli.” The Journal of General Physiology, vol. 77, no. 2, Feb. 1981, pp. 121-35.

[43] Norris, Noele. Mechanistic modeling of bacterial nutrient uptake strategies. (PhD thesis, Massachusetts Institute of Technology, 2019).

[44] Palsson, Bernhard . *Systems Biology: Constraint-Based Reconstruction and Analysis.* Cambridge University Press, 2015.

[45] Patzlaff, J. S., Heide, T. van der & Poolman, B. “The ATP/Substrate Stoichiometry of the ATP-binding Cassette (ABC) Transporter OpuA.” J. Biol. Chem. 278, 29546?29551 (2003).

[46] Peebo, K. et al. “Proteome reallocation in Escherichia coli with increasing specific growth rate.” Molecular BioSystems 11, 1184?1193 (2015).

[47] Poolman, B. et al. “Functional Analysis of Detergent?Solubilized and Membrane?Reconstituted ATP?Binding Cassette Transporters.” in Methods in Enzymology vol. 400 429?459 (Academic Press, 2005).

[48] Rees, D. C., Johnson, E. & Lewinson, O. “ABC transporters: the power to change.” Nature Reviews Molecular Cell Biology 10, 218?227 (2009).

[49] Romano, A. & Saler, M. H. “Evolution of the Bacterial Phosphoenolpyruvate:Sugar Phosphotransferase System.” in The Evolution of Metabolic function (ed. Mortlock, R.) (CRC Press, 1992).

[50] M.-T. Sung et al., “Crystal structure of the membrane-bound bifunctional transglycosylase PBP1b from Escherichia coli,” PNAS, vol. 106, no. 22, pp. 88248829, Jun. 2009.

[51] Szenk, Mariola, Ken A. Dill, and Adam M. R. de Graff. “Why Do Fast-Growing Bacteria Enter Overflow Metabolism? Testing the Membrane Real Estate Hypothesis.” *Cell Systems* 5, no. 2 (August 23, 2017): 95104.

[52] UniProt, “murC - UDP-N-acetylmuramate–L-alanine ligase - Escherichia coli (strain K12) - murC gene & protein.” [Online]. Available: http://www.uniprot.org/uniprot/P17952. [Accessed: 22-Jun-2017].

[53] C Wandersman & M Schwartz, “Mutations that alter the transport function of the LamB protein in Escherichia coli,” *J. Bacteriol.*, 151: 15?21, 1982.

[54] F. B. Wientjes, C. L. Woldringh, and N. Nanninga, “Amount of peptidoglycan in cell walls of gram-negative bacteria.,” *J. Bacteriol.*, vol. 173, no. 23, pp. 76847691, Dec. 1991.

[55] A. D. Wolfson, J. A. Pleiss, and O. C. Uhlenbeck, “A new assay for tRNA aminoacylation kinetics.,” RNA, vol. 4, no. 8, pp. 10191023, Aug. 1998.

### 7 Additional figures

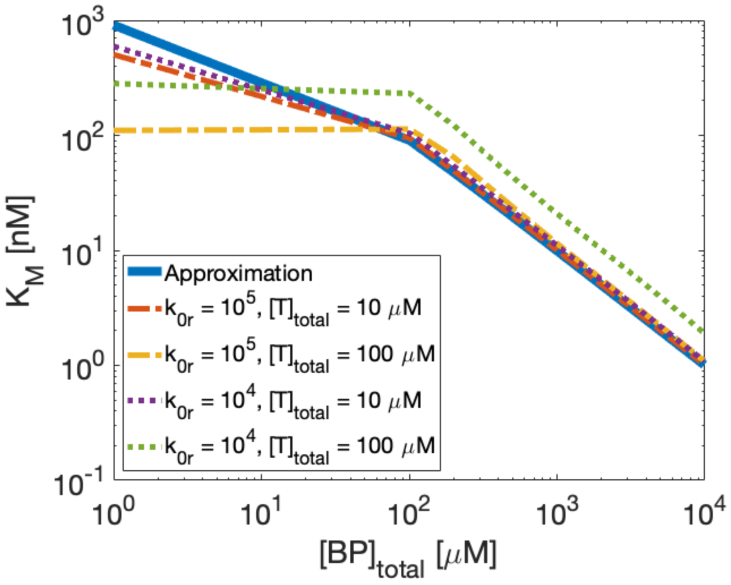

Figure S-6: **Comparison of approximate and exact ABC transport half-saturation concentration values.** Here we compare our Michaelis-Menten approximation of the half-saturation concentration for ABC transport with the exact half-saturation concentration obtained by solving the set of four equations for ABC transport rates using different values for the total abundance of transport units, binding association rate $k_{0r}$, and binding protein abundances ($x$-axis).

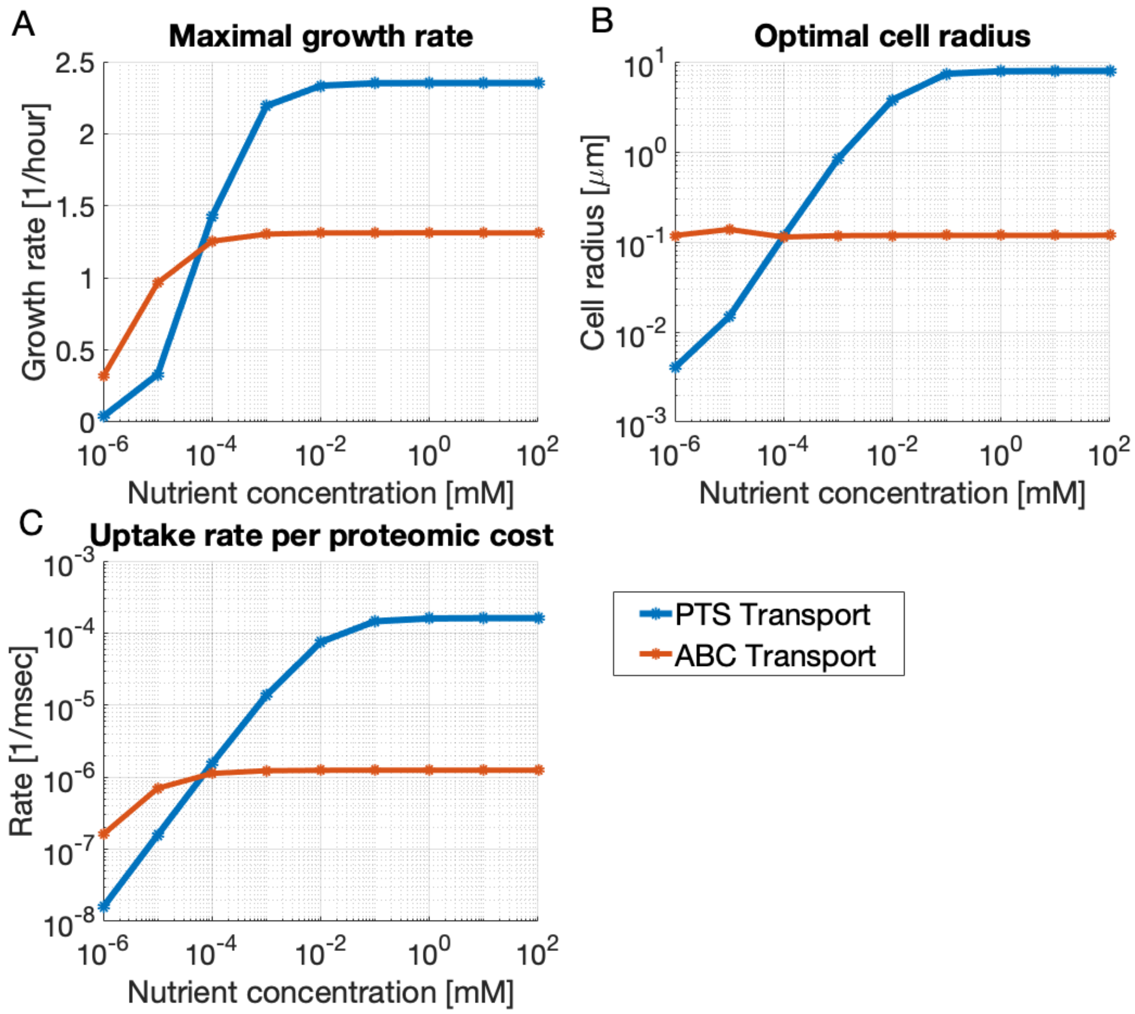

Figure S-7: **A rate-affinity trade-off.** These plots show the maximal growth rate obtained for PTS and ABC transport systems and their corresponding optimal cell radii (inversely related to the optimal surface-area-to-volume ratio) and uptake rate per proteomic cost. ABC transport achieves higher growth rates at low nutrient concentrations because it can achieve higher specific affinities per proteomic cost. Conversely, PTS transport achieves higher growth rates at high nutrient concentrations because it can achieve higher maximal uptake rates per proteomic cost.

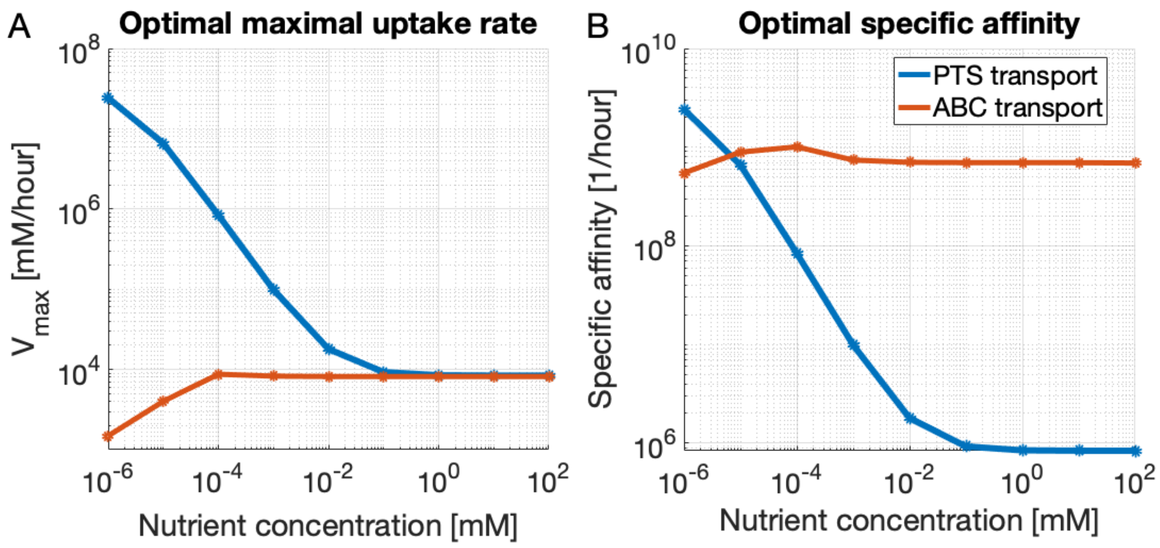

Figure S-8: **Optimal maximal uptake rates and specific affinities.** Here are plots showing the corresponding optimal maximal uptake rates ($V_{\text{max}}$) and optimal specific affinities ($V_{\text{max}}/K_{\text{M}}$). Although, in this model, PTS transport has a higher optimal specific affinity than ABC transport at 1nM, achieving this specific affinity has a much higher proteomic cost because it requires expressing more transport units, thereby also increasing the maximal uptake rate.

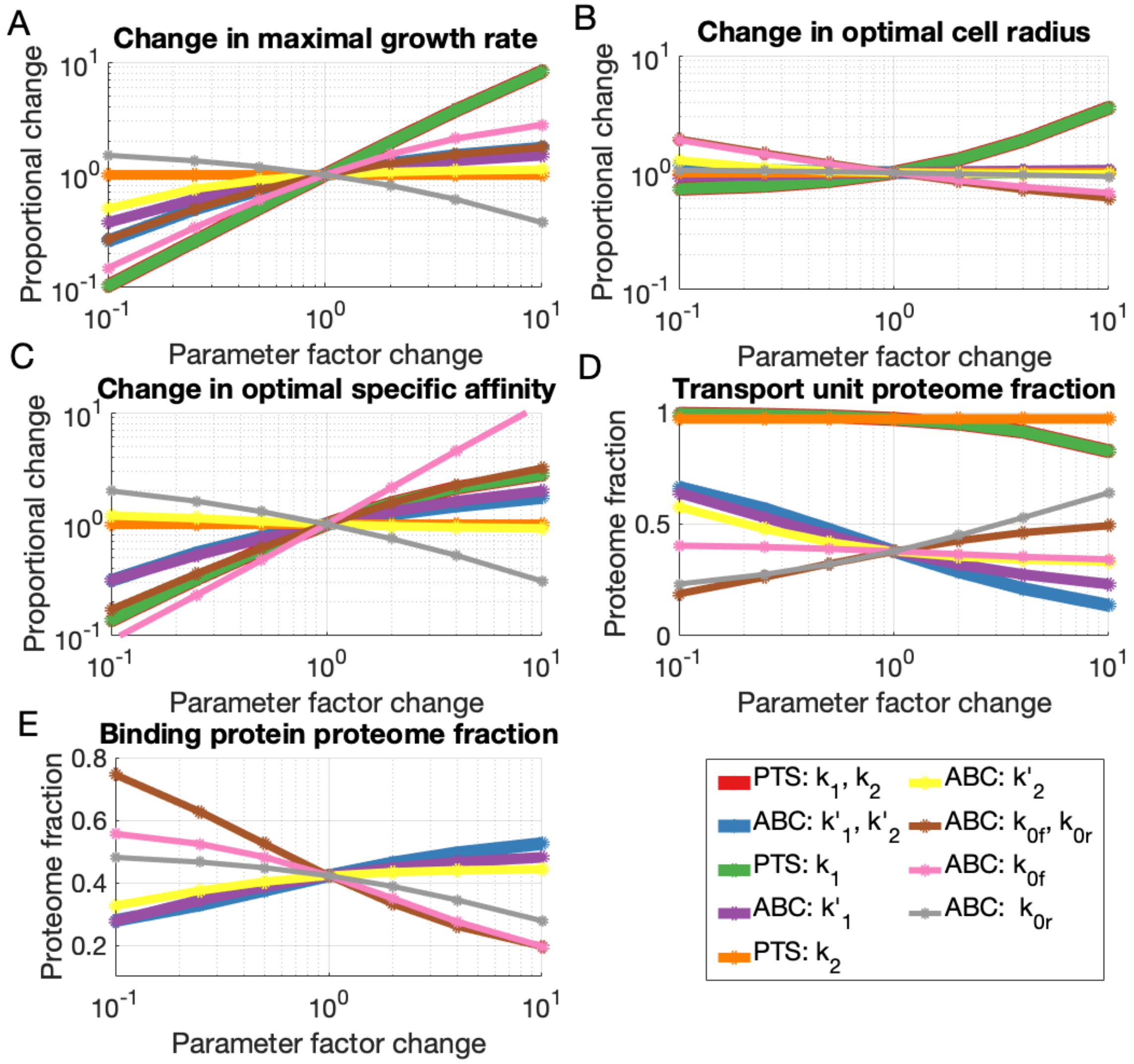

Figure S-9: **Sensitivity analysis of PTS and ABC transport systems at $[\mathrm{S}]_{\mathrm{ext}} = 1$ nM over various transport kinetics rates.** To ensure that the observed general trends did not depend on the transport kinetic rates, we conducted a suite of sensitivity analyses in which we modified the specified transport kinetic rate by a factor change ($x$-axis).

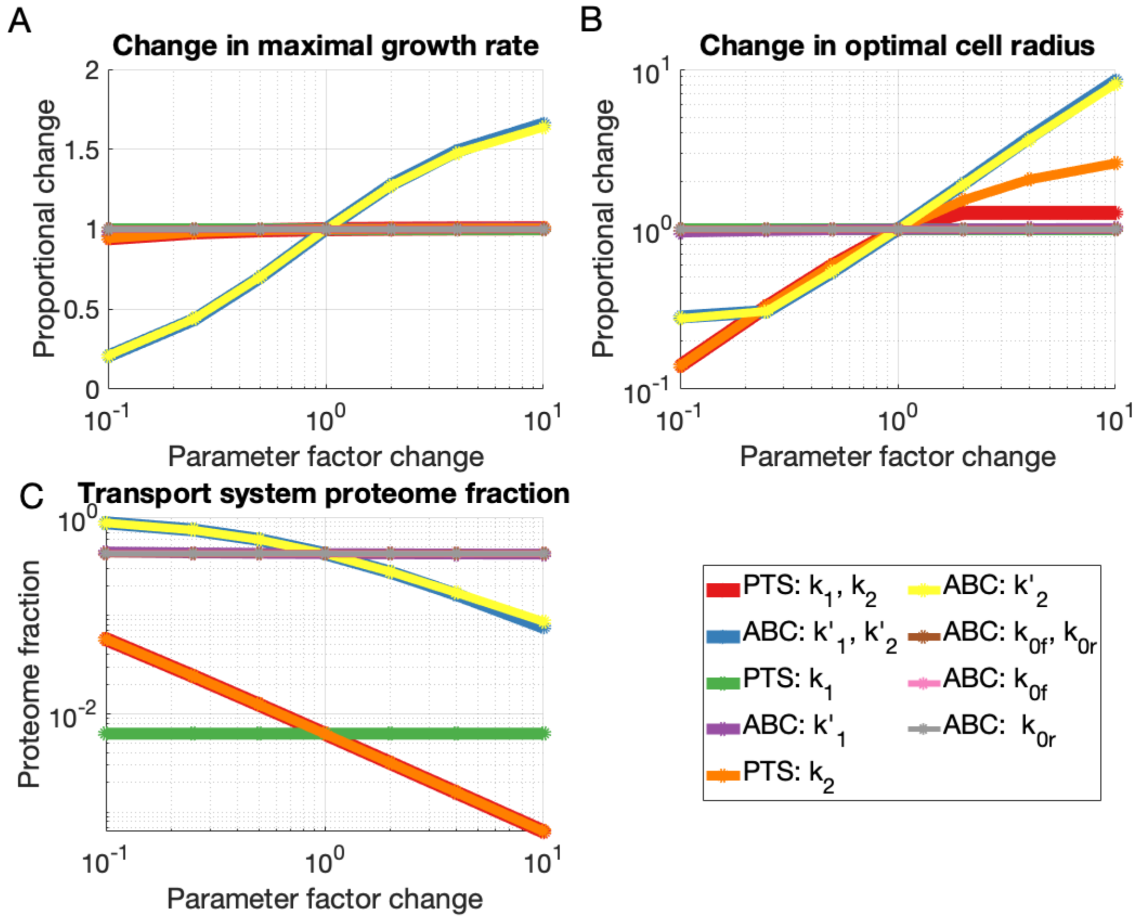

Figure S-10: **Sensitivity analysis of PTS and ABC transport systems at saturation over various transport kinetics rates.** To ensure that the observed general trends did not depend on the transport kinetic rates, we conducted a suite of sensitivity analyses in which we modified the specified transport kinetic rate by a factor change ($x$-axis).

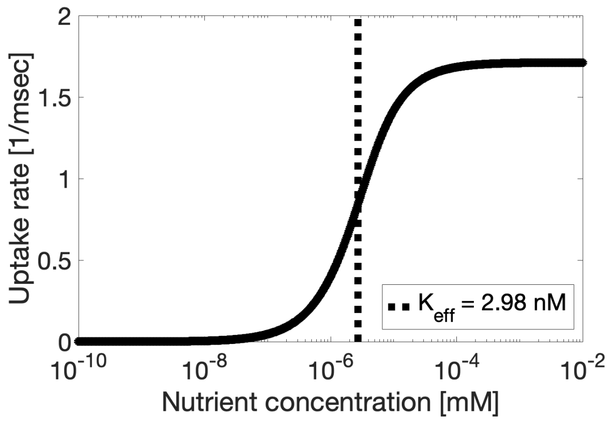

Figure S-11: **Calculating the half-saturation concentration of ABC transport.** To calculate the effective half-saturation concentration of an optimal solution to a particular proteome allocation problem, we used the set of four equations describing ABC transport to determine the uptake rate over a range of nutrient concentations ($x$-axis). Here we show the calculated uptake rates over various nutrient concentrations for the proteome allocation obtained when optimized the cell for growth at an extracellular concentraiton of $[S]_{\mathrm{ext}} = 1$ nM.

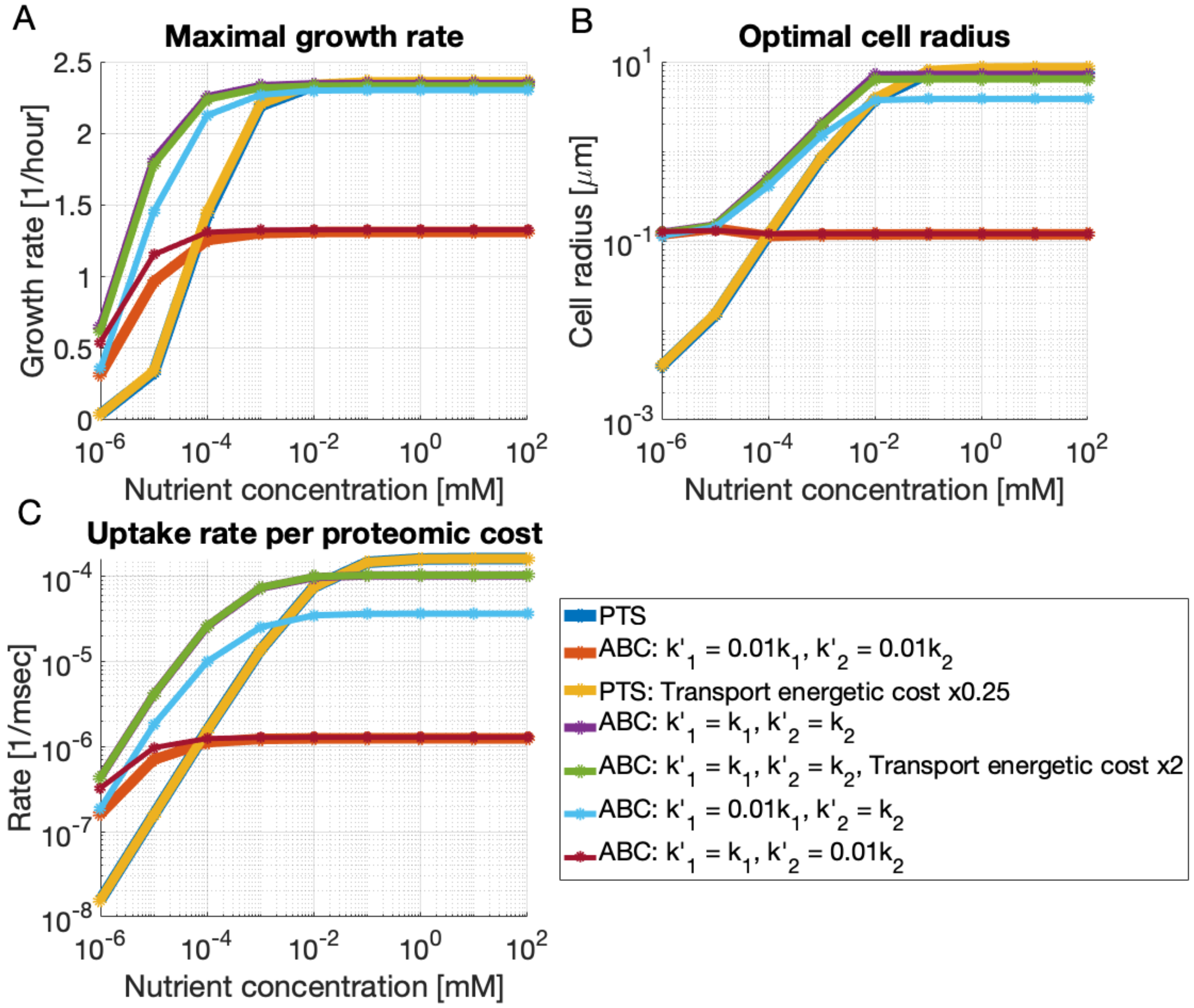

Figure S-12: **Sensitivity analysis of the rate-affinity trade-off.** These plots show results of proteome allocation problems with variations in the PTS and ABC transport systems. The proteome allocation problem is solved for different extracellular substrate concentrations (x-axis). (A) shows the maximal growth rates achieved with the optimal proteome allocation, (B) shows the optimal cell radii used to achieve those maximal growth rates, and (C) shows the uptake rate per proteomic cost, which is the rate of carbon consumption divided by the amount of carbon comprising the porins, binding proteins, and membrane-bound transport units. We argue that ABC transport rates, $k_1'$ and $k_2'$, are likely on the order of one hundred times lower than PTS transport rates because of the slow diffusivity of the bulky binding proteins and because, when ABC transport has the same transport rates as PTS transport (purple), they outcompete PTS transport at all concentrations. PTS transport systems may have lower energetic costs (yellow) and ABC transport rates may have higher energetic costs (green), but these modifications do not considerably alter maximal growth rates at high nutrient concentrations. The lower rates of ABC transport (red, maroon) result in smaller optimal cell radii.

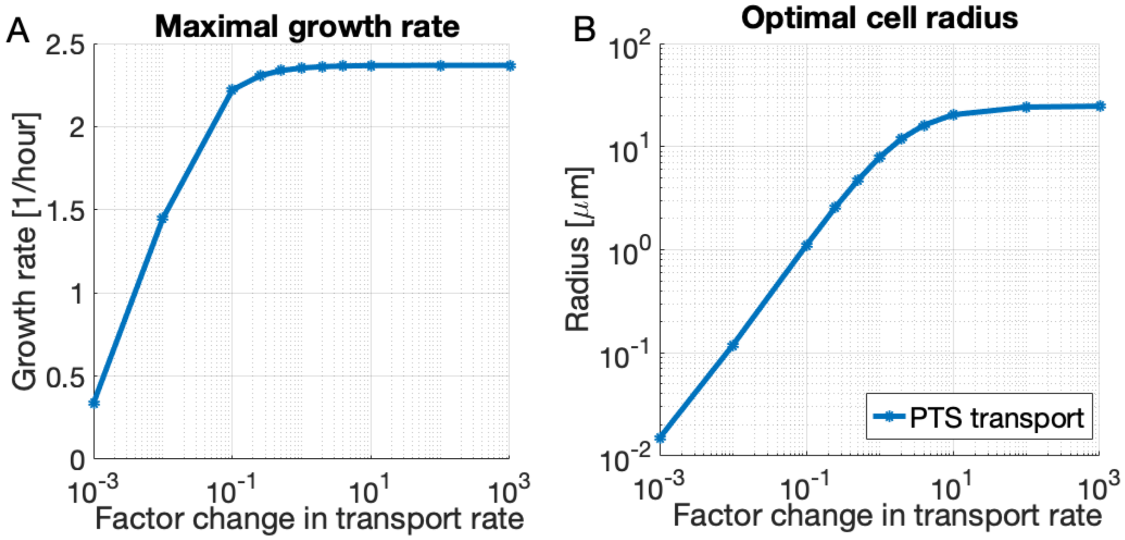

Figure S-13: **Sensitivity analysis of proteome allocation for PTS transport to changes in translocation rate $k_2$ at extracellular carbon concentration $[S]_{\mathbf{ext}}$ = 100 mM.** Increases in the translocation rate result in (A) higher achievable growth rates and (B) larger optimal cell radii (that is, smaller surface-area-to-volume ratios).

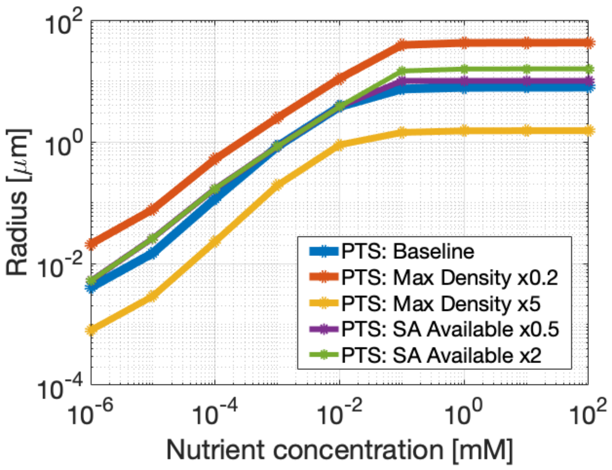

Figure S-14: **Active constraints on cell radius.** Both the surface area "real estate" constraints and the density constraints are active for the PTS transport proteome allocation problem. Increases in maximal allowed density result in smaller optimal cell radii (red and yellow). Increases in the fraction of the surface area available to the membrane-bound transport units result in larger optimal cell radii (purple and green).

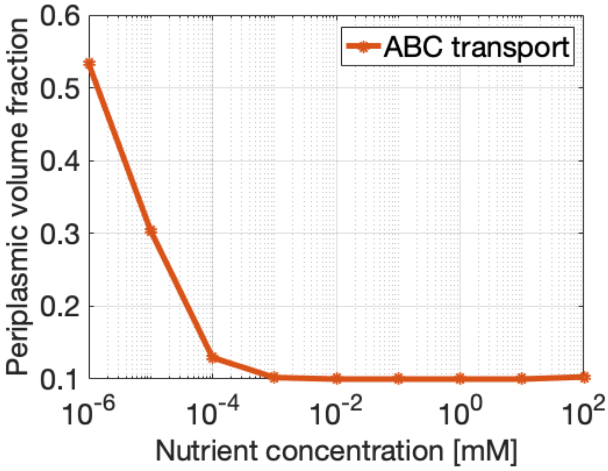

Figure S-15: **Optimal periplasmic volume fraction for ABC transport.** The optimal periplasmic volume fraction increases as nutrient concentration decreases to allow for greater abundances of binding proteins, which are subject to a density constraint on the periplasm.
